# Evolutionarily conserved neural dynamics across mice, monkeys, and humans

**DOI:** 10.64898/2026.03.06.709637

**Authors:** Olivier Codol, Margaux Asclipe, Anton R. Sobinov, Zihao Chen, Junchol Park, Nicholas G. Hatsopoulos, Joshua T. Dudman, Juan A. Gallego, Guillaume Lajoie, Matthew G. Perich

**Author notes:** co-first authors (Olivier Codol, Margaux Asclipe).

## Abstract

On evolutionary timescales, brain circuits adapt to support survival in each species’ ecological niche. While some anatomical aspects of neural circuitry are conserved across species with distant evolutionary origins, each species also exhibits specific circuit adaptations that enable its behavioral repertoire. It remains unclear whether homologous brain regions leverage analogous neural computations as different species perform common behaviors such as reaching and manipulating objects. Here, we directly assessed conservation of neural computations using intracortical recordings from mouse, monkey, and human motor cortex—a homologous region across many mammals—during motor behaviors crucial for survival. We hypothesized that, despite their phylogenetic distance, rodents and primates produce movements through conserved neural computations implemented by motor cortical population dynamics. Remarkably, we found that movement-related neural dynamics were highly conserved across species, while variations in behavioral output were uniquely captured in neural trajectory geometries. Strikingly, neural dynamics during movement across species were more conserved than those across brain regions in the same human and between motor preparation and execution in the same monkeys. Lastly, through manipulation of neural network models trained to perform reaching movements, we reinforce that conservation of neural dynamics across species likely stems from shared circuit constraints. We thus assert that evolution maintains neural computations across phylogeny even as behavioral repertoires expand.

## Introduction

Over millions of years, natural selection and other pressures guide changes in the behavioral capabilities of each species^1,2^, yielding behaviors tailored to their ecological niches (**Fig. 1a**). To achieve these behaviors, brains similarly evolved in complexity, from their early origins as small networks of cells implementing primitive stimulus-response computations^2,3^, to the emergence of functionally and anatomically specified, recurrently-connected brain regions^4^. The neural circuits in these regions were shaped over long timespans as species evolved and diverged by building on existing structures through a process of phylogenetic refinement^2^. Consequently, across mammalian species, the wide array of brain regions that guide behavior often have common evolutionary ancestry. For example, the motor cortex is recognizable (based on cellular analysis^5^ and anatomy^6^) in many mammalian species^7–12^. There is considerable evidence for cellular, anatomical, and organizational similarities in cortical and subcortical regions across diverse species^5,11,13–15^. Yet, even across mammals such as mice and primates there are numerous notable differences^16,17^ such as monosynaptic corticospinal connections^18^ and a wider range of cell subtypes^5^ in primates. These variations from shared evolutionary ancestors are associated with adaptations to the ecological niche of each species. Consequently, it remains unclear whether the *neural computations* that homologous cortical brain regions use to produce shared behaviors are conserved throughout evolution, or whether new behavioral adaptations alter the computational substrate of the cortex. Uncovering whether convergent or shared neural computations exist throughout the animal kingdom is critical to refining our theories of the biological implementation of processes that regulate animal behavior.

**Figure 1.**
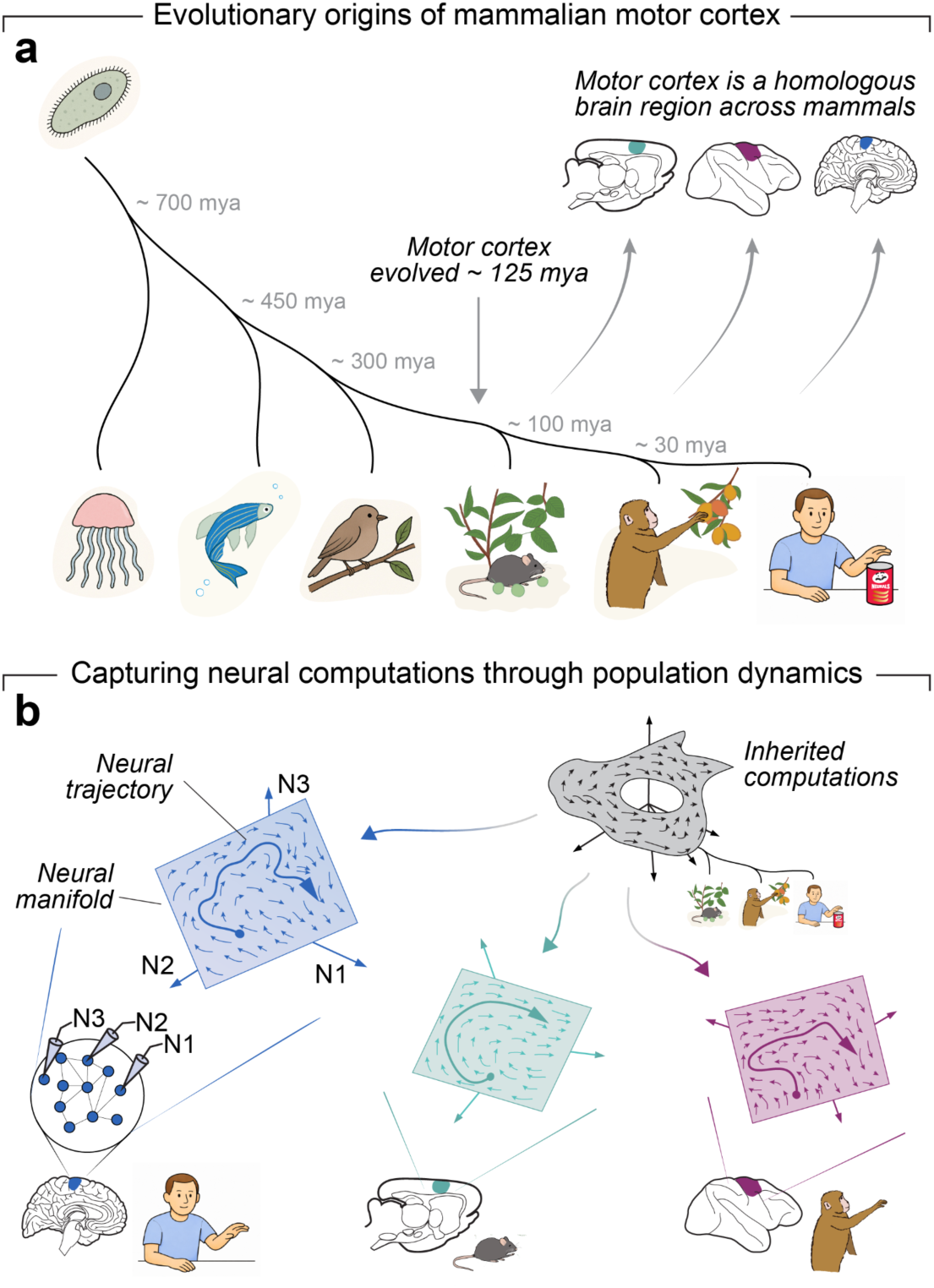
Hypothesis of evolutionarily conserved neural computations through population dynamics. (a) Over hundreds of millions of years, the animal kingdom evolved a spectrum of animals with behavioral repertoires adapted to their environmental niche. The emergence of the motor cortex approximately 125 million years ago (mya) aided many modern species in acquiring dexterous arm/forelimb behaviors such as grasping and manipulating objects. We explore whether neural population activity in mice, monkeys, and humans share similar computations, quantified by neural dynamics and geometry. (b) Neural computations are implemented through dynamical “rules” that constrain neural activity during behavior (referred to here as the *dynamics*), which can often be visualized as a “flow field” in a neural state space capturing the activity of recorded neurons (N1-3 for the three neuron example here). By comparing these neural population dynamics across species such as mice, monkeys, and humans in homologous regions such as the motor cortex, we can directly test whether neural computations are conserved throughout evolution.

To tackle this open question, we focus on a motor behavior that is nearly ubiquitous across the animal kingdom: engaging forelimbs to reach for, grasp, and manipulate objects. Mammalian species from rodents to primates engage forelimbs to achieve these behaviors under the guidance of the motor cortex^5^, a region with ancient evolutionary origins (on the order of 125 million years ago^11^) and a direct role in driving movements^7,11^. We directly explored conservation of neural computations across species in the motor cortex using the computation through dynamics framework^19^. This framework states that the temporal progression of neural activity along a neural manifold^20^ is dependent on both past activity and inputs to the population, subject to biophysical constraints such as recurrent connectivity profiles^19,20^. These dynamics underlie the neural population activity that we can observe as individuals move and interact with the world. In this light, populations of neurons enacting the same neural computations should also exhibit similar dynamics during the production of behavior^21^. The temporal evolution of neural dynamics can be described with *trajectories* along the neural manifold, which trace activity in a state space defined by the firing rate of each constituent neuron^20,22^ (**Fig. 1b**). These trajectories are subject to underlying rules that govern their unfolding over time^19^ (which we refer to as the dynamics for succinctness), often visualized as flow fields in a neural state space in which each axis represents the activity of a recorded neuron. When considering two neural trajectories, such as those from two different species, quantitative comparison of these flow fields becomes a direct proxy for the similarity of the underlying neural computations^23^.

We hypothesized that, if the motor cortex operates by conserved computations across evolutionarily divergent species, we should see conservation of the dynamics of neural population activity across species during similar motor behaviors. We tested this hypothesis directly using intracortical neural population recordings from the motor cortex of mice^24^, macaque monkeys^25,26^, and a human participant with cervical spinal cord injury who was enrolled in a clinical trial^27^ as they performed several skilled arm/forelimb motor behaviors focused on object interactions. We found a striking conservation of neural dynamics across all three species, even in the face of considerable variability in the nature of the motor behaviors the individuals performed. This demonstrates that even across species the motor cortical activity unfolded with conserved “dynamical rules”. To underscore this conservation, we show that neural dynamics change when behavioral demands require different neural computations: (1) between motor and somatosensory cortex (brain regions that both contribute to dexterous motor control^7^) in humans; and (2) between motor preparation and execution in macaques. We then demonstrate that differences in motor output across both individuals and species are captured in the specific geometry (or shape) of trajectories produced by the motor cortex for the given behavior, allowing one conserved dynamical system to produce a range of motor outputs based on each species niche. Finally, we underscore that circuit constraints are intimately tied to dynamical similarity using architectural manipulations to recurrent neural network models trained to move a biomechanical simulation of a primate arm. We thus posit that as evolution shaped the genomes specifying the brains of mice, monkeys, and humans, neural computations were retained and repurposed as behavioral repertoires expanded. Our findings have implications for the translation of fundamental studies in animals to guide our understanding of human brain function and treatments for brain disorders.

## Results

### A multi-species dataset to study neural computations in the motor cortex

To explore neural dynamics across species, we compiled a dataset of intracortical neurophysiology recordings during motor behaviors in mice, macaque monkeys, and humans. Four mice were trained to perform a lever pull task^24^ (**Fig. 2a**). In this task, mice waited for a cue before reaching for, grasping, and pulling on a lever. The lever was placed in two possible positions and loaded with two different masses. We also studied a cohort of four monkeys trained to perform a highly similar task where they reached for and pulled on objects placed in the coronal plane in front of them (**Fig. 2b**). These were pulled from two different datasets^25,26,28,29^. In the first dataset, two monkeys used a power grasp to pull on a horizontally-oriented cylinder placed in eight different positions^25^. In the second dataset, two monkeys grasped and pulled on a cubic object in one position^26^. In this dataset, the monkeys used two different grip types (side grip or precision grip) and pulled on two different loads (low and high). Thus, there was considerable variability in the object locations, object shapes, grip types, and required pulling forces, giving a richer set of behavioral states in monkeys compared to the mouse tasks. We compared the mouse and monkey data with recordings from the motor cortex of a human participant in a clinical trial^27^. The participant had an incomplete spinal cord injury (level C4) but was able to reach for and transport an object using a modified grasp strategy that involved wrist extension with minimal active motor function of the intrinsic or extrinsic hand muscles. We recorded four sessions where the participant reached for, picked up, and moved a cylindrical object after receiving an auditory and visual cue (**Fig. 2c**).

**Figure 2.**
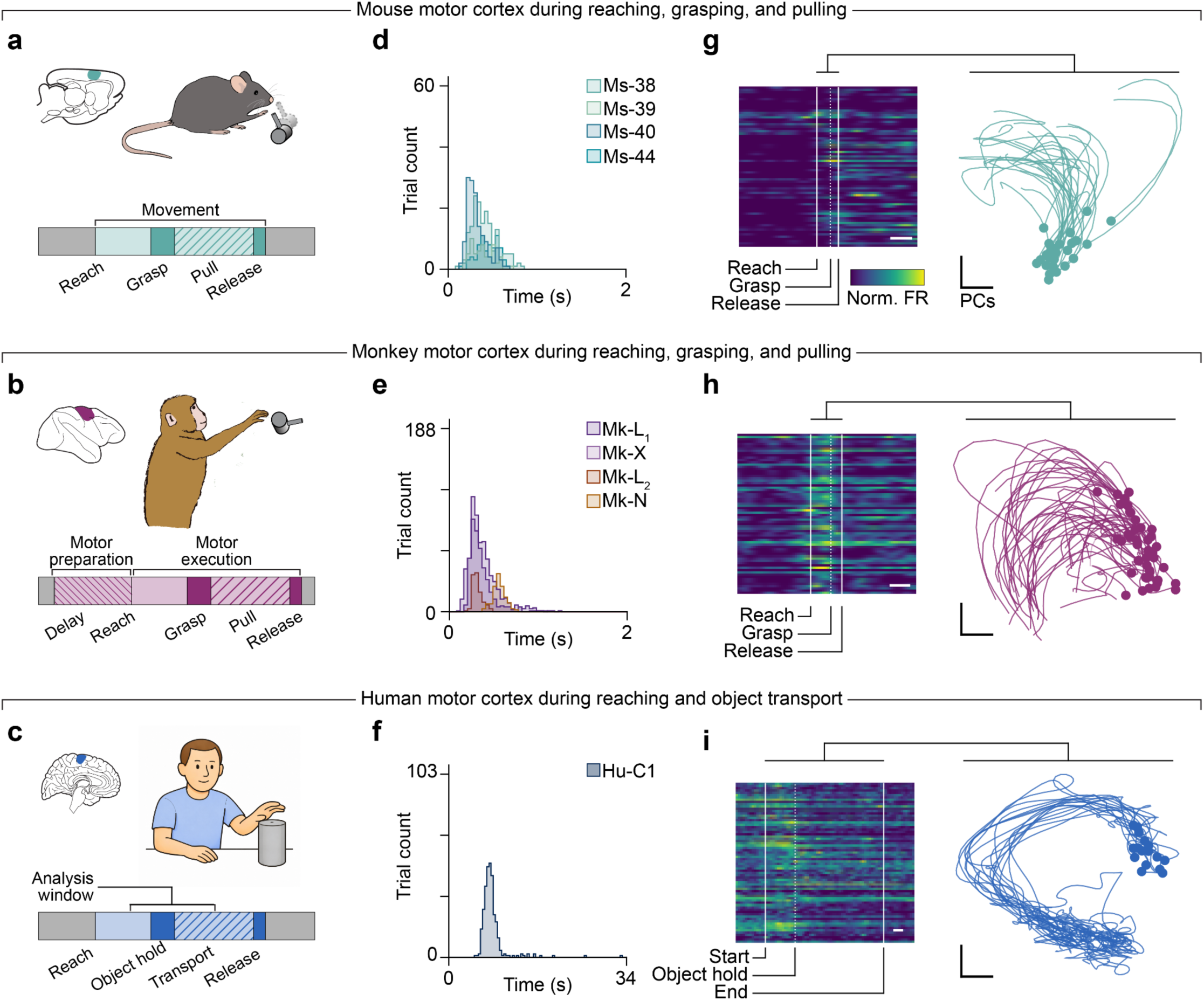
A multi-species motor neurophysiology dataset. (a) Overview of the mouse lever pull task. The mice reached for, grasped, and pulled on the lever to receive a reward. The mice self-initiated the movement after the lever appeared in one of two locations, and the lever had two force profiles. (b) Overview of the monkey object pull task. The movements were similar to the mouse task, involving reaching, grasping objects, and pulling on them. Across datasets, the monkeys reached for differently shaped objects in multiple locations. Unlike the other species, the monkeys performing this task were trained to accommodate a delay period to enable motor planning before the go cue. (c) Overview of the human object transport task. This task involved picking up an object and moving it to a cued location on a table. The resulting movements were more variable across trials than the well-trained monkeys and mice. (d-f) Histograms of the trial durations for all pooled across all sessions for each mouse, monkey, and human. **(g)** Left: heatmap of neural activity for one trial from Ms-40. Colorbar shows normalized activity scaled per channel. Solid lines indicate the start and end of the movement analysis window, and dashed line indicate the time of object acquisition. White scale bar indicates 200 ms. Right: example neural trajectories for 28 representative trials from the same Ms-40 session. Circles indicate the movement onset. The thicker line indicates the example trial shown in Panel c. **(h)** Example neural activity from Mk-L in the cylindrical object dataset. Data presented as in Panel g. **(i)** Example neural activity from one human session in the object transport task. Data presented as in Panel g.

The behaviors across all species involved analogous reaching and grasping movements, yet with variation in specific behavioral features such as reaching distance, grip orientation, or even effector (i.e., monkey arm vs mouse arm). The differences were especially apparent in the dataset. When we quantified the time required to complete the task, we found that it was identical for nearly all mice and monkeys (**Fig. 2d,e**) with the exception of one monkey from the second (cubic object) dataset who reached significantly slower but spent similar time pulling (**Fig. S1b**). Due to the nature of the less structured human object transport task, trials took considerably longer to complete (**Fig. 2f**). To enable fair comparison to the animal data, we isolated a shorter window of time surrounding object acquisition (see Methods). As the individuals of each species performed their respective behaviors, neural populations were recorded from the primary motor cortex of mice using Neuropixels probes^30^ (**Figs. 2g**, **S2a**), monkeys using various chronically-implanted microelectrode arrays (**Figs. 2h**, **S2b,c**), and the human participant using NeuroPort Electrode (Blackrock Neurotech) microelectrode arrays^31^ (**Figs 2i**, **S2d**). To visualize the neural population activity across species, we applied Principal Component Analysis (PCA) to reduce each session’s population activity to a 20-dimensional space, which explained the majority of variance across all datasets (**Fig. S3**). Motor cortex in mice and monkeys showed smooth, relatively simple trajectories, tracing out a single path throughout the reaching, grasping, and pulling phases (**Figs. 2g,h**, **S2a-c**). The human motor cortex (**Figs. 2i**, **S2d**) was more complex, especially during object manipulation, with occasional loops and divergences. In the subsequent analyses, we explore whether these trajectories represent conserved neural computations across the three species.

### Quantifying dynamical similarity of neural activity across datasets

We tested our hypothesis that neural computations are conserved throughout evolution using Dynamical Similarity Analysis (DSA)^23^ (**Fig. 3a**) applied to the motor cortical population activity^19^. DSA allows us to directly measure the similarity of the dynamics underlying the observed neural trajectories across individuals both within and across species. In brief, DSA transforms the (presumably nonlinear^32^) neural dynamics into a new, linearized state space before fitting dynamical systems models capturing the temporal unfolding of the neural trajectories (see Methods and Ref. 23 for more details). From these models, we obtain a state transition matrix, ***K***, for each dataset which captures the “rules” by which the system evolves over time. The dynamical similarity can then be easily quantified by computing a distance between the transition matrices ***K*** of each dataset and learning an optimal alignment, ***C***, between the two matrices. We performed extensive validation to ensure we selected adequate hyperparameters for the model fits and obtained similar reconstructions of neural dynamics for each dataset for a range of hyperparameters and assumed neural dimensionalities (**Fig. S4**, see Methods). When applied to our multi-species motor cortical dataset (**Fig. 2**), DSA learned transition matrices ***K*** which captured the vast majority of variance in the neural trajectories for all sessions (**Fig. 3b-c**, **S5**).

**Figure 3.**
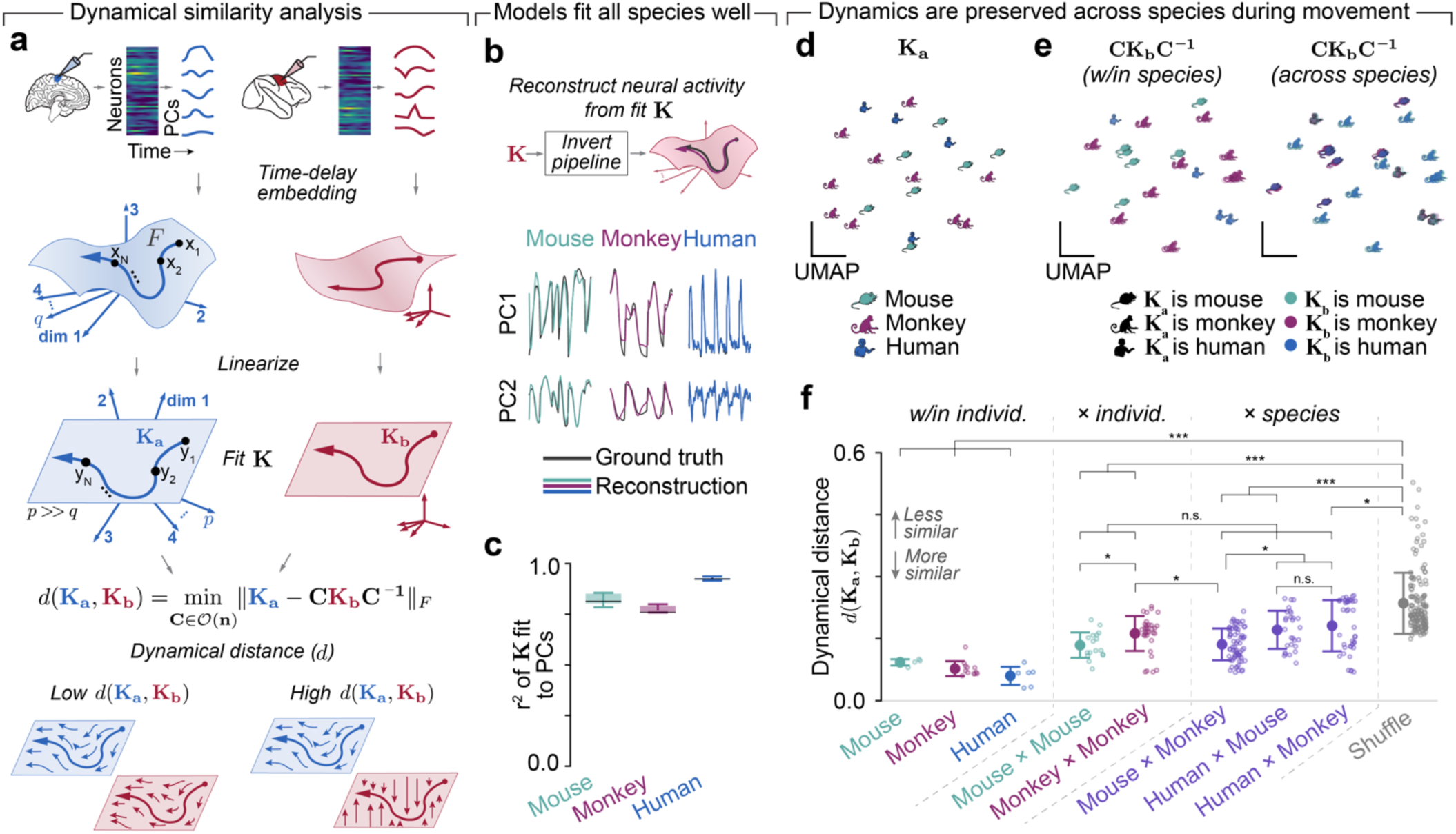
Dynamical similarity across species of neural population recordings in the motor cortex. (a) Dynamical similarity analysis (DSA) compares two sets of neural trajectories (*a* and *b*) that are transformed by time-delay embedding (F). The result metric represents the distance (*d*) after aligning linearized dynamical systems models fit to each dataset (K_a_ and K_b_) via a linear transformation C. (b) The leading two motor cortical PCs (black lines) for five example trials over time from the sessions visualized in **Fig. 2d-f** and reconstructed from dynamical systems fits from **K** for that session for mice (green, left), monkeys (red, middle), and humans (blue, right). PC1 and PC2 are proportionally scaled based on the relative variance explained. (c) Distributions of R^2^ values for transition matrices K fit to each session for all species. Box plot: black line shows median, shaded area shows quartile, whiskers show 95% range (mice, N=147; monkeys, N=210; humans, N=84). (d) UMAP embedding visualization of all K_a_ matrices fit to motor cortical recordings for all sessions of each species. (e) UMAP visualization of all aligned **K_b_** matrices. Left: the subset of session pairs where both **K_a_** and **K_b_** are fit to the same species. Right: the subset of session pairs where **K_a_** and **K_b_** are fit to different species. The shape shows the target species (**K_a_**) and the color shows the aligned species (**K_b_**). Session pairs cluster within the embedding according to **K_a_**. Note that the left and right panels represent the same UMAP embedding computed on both sets of points, and that the UMAP embedding of Panel d was performed independently. (f) Dynamical distance values found by DSA within individuals in mice (green, N=4), monkeys (red, N=12), and the human participant (blue, N=6). We also show distance across individuals in mice (N=17) and monkeys (N=33). Since we had only one human participant, we could not compare variability across individuals. Both plots show the same embedding space just separated by type for visualization given the substantial overlap based on **K_a_**. These are reference values for our across species comparisons (purple): mouse and monkey (N=70), human and mouse (N=28), and human and monkey (N=40). We computed a control distribution found through shuffling chunks of data (see Methods) as an “upper bound” (gray, N=153). Error bars represent mean ± s.e.m., with individual datapoints plotted beside them. All statistical comparisons: Wilcoxon’s rank-sum test; *: *p*<0.01; **: *p*<0.001; ***: *p*<0.0001; n.s.: not significant. For visual clarity given the large number of groups, we present only the most relevant statistical comparisons on the panel. See **Table S1** for the full set of exact p-values for all comparisons.

### Neural dynamics are conserved across species in a homologous brain region

We next used the transition matrices ***K*** for each recording to test the hypothesis that neural dynamics are conserved across species in the motor cortex. As a first exploration, we visualized the fit ***K***s with a two-dimensional embedding from UMAP^33^ (**Fig. 3d**). We found no systematic separation across species. Interestingly, the aligned ***K***s (**Fig. 3e**) also clustered together, indicating similar dynamical systems model fits. Comparison of the learned transition matrices ***K*** with DSA produces a distance metric reflecting the dissimilarity of the dynamical system underlying the neural population activity. Since the absolute value of this metric can be difficult to interpret, we established two key reference values. First, as an upper-bound, we computed a null distribution from a shuffle control (“Shuffled”; see Methods). Second, as a lower bound, we compared neural dynamics across sessions within each individual, representing recordings from the same brain. We anticipated that any across-species comparisons should lie within these two extremes. As our most important reference, we established the similarity of neural activity *within* species (“x individual”) by applying DSA across sessions from different mice or monkeys (note that since we obtained data from a single human participant, we were unable to quantify variability across human individuals). Here, we make the reasonable assumption that similar neural computations underlie movement in different individuals^21^, providing a direct expectation for what dynamical distance values to expect if different species generate movements using conserved neural computations.

We finally applied DSA to directly compare across all permutations of species: mouse to monkey, human to mouse, and human to monkey (**Fig. 3f**, right). Remarkably, we found that each pair of species were as similar, dynamically, as any two individuals of a single species. To support this conclusion, we performed several further analyses. First, the similarity across mice and monkeys could in principle be dominated by a single phase of the structured behavioral task, yet we observed high dynamical similarity across species even when separately comparing the reaching and pulling phases independently (**Fig. S6a**). Second, we confirmed that our assumed neural dimensionality of 20 PCs did not bias our conclusions: we found dynamical similarity across species for a range of different dimensionalities (**Fig. S6c**). Third, we demonstrated that dynamical similarity across species did not depend on our choice of alignment window within the longer human trials: we saw equivalent dynamical similarity with shorter windows misaligned to the time of object acquisition (**Fig. S6d,e**). Lastly, we confirmed that the dynamical distance from DSA matches our intuitions about the generalization of the dynamics models across brains.

In summary, our findings and control analyses identify high dynamical similarity in motor cortical activity between mice, monkeys, and humans. This is particularly surprising in light of the known biomechanical and neuroanatomical differences between rodents and primates, such as more direct corticospinal control of motoneurons between mice and monkeys^18^, or the apparent reduction of rubrospinal pathways between monkeys and humans^34^.

### Neural dynamics are not conserved between motor and somatosensory cortex

If similarity in neural dynamics is due to conserved neural computations in the motor cortex, our analysis should identify dissimilar neural dynamics if the computations implemented by a neural population are different^19^, as we should expect in functionally distinct brain regions^4^. We thus assessed dynamical similarity across anatomically and functionally distinct brain regions in the same individual. The human participant was simultaneously implanted with a second set of electrode arrays on the postcentral gyrus in the cutaneous primary somatosensory cortex^27^, allowing us to study cortical responses to sensation during movement. Since primary sensory regions are strongly driven by peripheral inputs—unlike the motor cortex, which is primarily fed by recurrent inputs from other brain regions^7^—we predicted that these two regions would implement different computations, evidenced by different dynamics (**Fig. 4a**). Indeed, we saw strikingly different population activity in the neural activity of somatosensory cortex (**Fig. 4b, S2d-e**) compared the activity previously studied for the motor cortex (**Fig. 2i**). Yet, even within the same session during the same behavior, their dynamics were significantly dissimilar by DSA and comparable to values in our control distribution. When we compared each region to itself across sessions with DSA, we found equivalent self-similarity of neural dynamics for both motor and somatosensory cortex (**Fig. 4c**). Altogether, these analyses underscore that neural dynamics are not conserved after changes in the computations implemented by a neural population.

**Figure 4.**
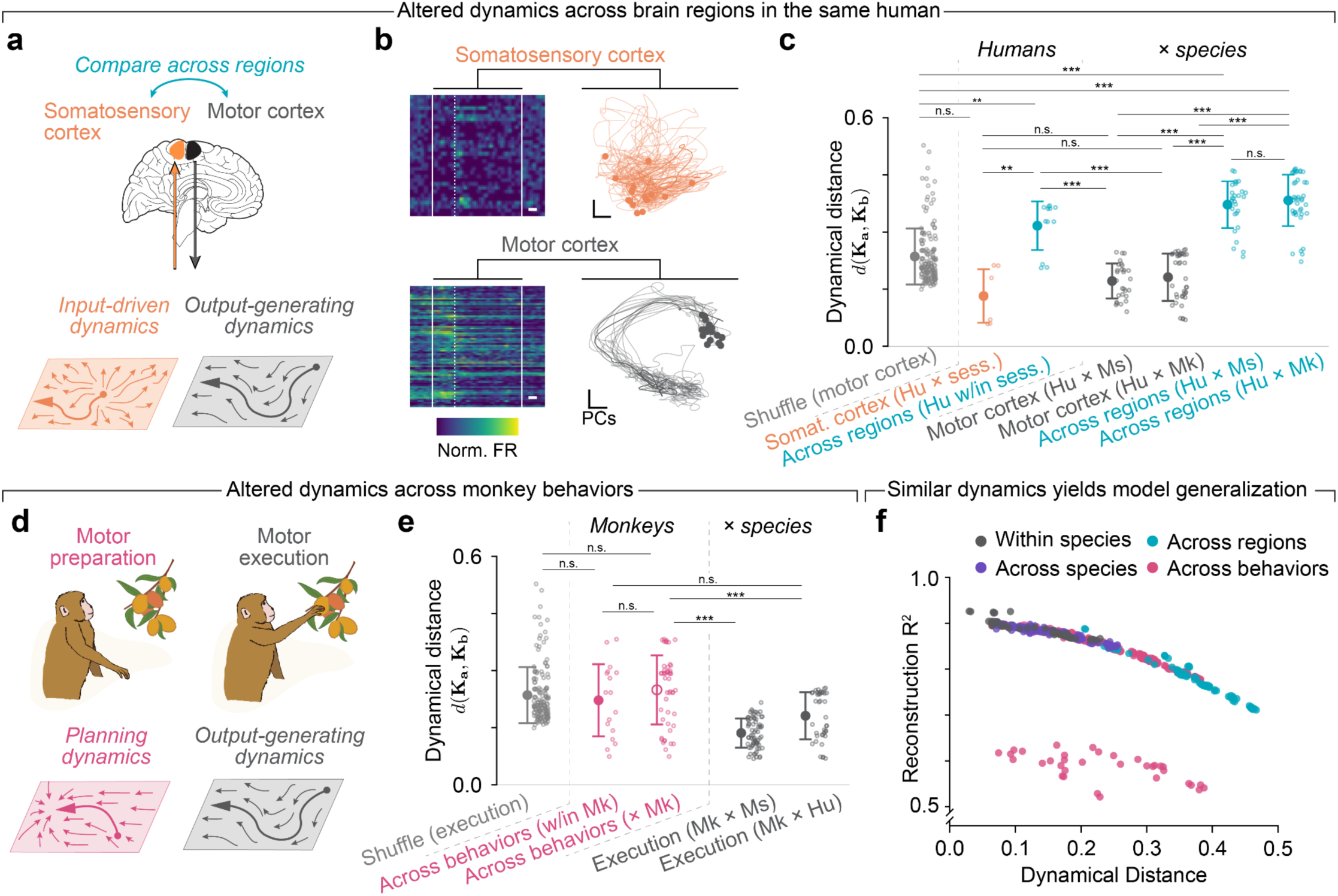
Neural dynamics are not conserved in cases where neural computations change. (a) Motor and somatosensory cortex in humans are driven by distinct inputs for different purposes: the generation of robust motor output and processing of somatic feedback, respectively. (b) Example activity comparing somatosensory (top) and motor (bottom) cortex in the human participant. Data presented as in **Fig. 2g**. Right: example neural trajectories for human motor cortex as in **Fig. 2g**. These neurons were recorded simultaneously with the somatosensory cortical neurons. (c) Dynamical distance across sessions for somatosensory cortex (orange), and across regions on the same sessions with simultaneous multi-regional recordings (teal). Data presented as in Panel b. The across species (dark gray) and shuffle (light gray) distributions are reproduced from **Fig. 3f**. All statistical comparisons: Wilcoxon’s rank-sum test; *: *p*<0.01; **: *p*<0.001; ***: *p*<0.0001; n.s.: not significant. For visual clarity given the large number of groups, we present only the most relevant statistical comparisons on the panel. See **Table S1** for the full set of exact p-values for all comparisons. (d) Motor preparation and execution likely involve distinct computations with divergent inputs and immediate outputs. (e) Dynamical distance across states (magenta) both within session (w/in, filled circle) and across sessions (hollow circle). Within session comparisons represent the same neurons on the exact same trials in the same monkey. Data presented as in Panel b. All statistical comparisons: Wilcoxon’s rank-sum test; *: *p*<0.01; **: *p*<0.001; ***: *p*<0.0001; n.s.: not significant. For visual clarity given the large number of groups, we indicate only the most relevant statistical comparisons. See **Table S1** for the full set of exact p-values for all comparisons. (f) Scatter plot relating the reconstruction performance in the linearized dynamics space of the aligned K_b_ models across sessions to the dynamical distance for that pair of sessions.

### Neural dynamics rapidly change to reflect computational needs across behaviors

Our hypothesis also makes the prediction that neural dynamics should be dissimilar when behavioral demands are different, changing the required neural computations. Prior work has demonstrated the preparation and execution of movement should be considered as two distinct computations^35,36^ (**Fig. 4d**). We thus hypothesized that neural dynamics would not be conserved across these two computations even within species, meaning that differences between preparation and execution within a species should be greater than differences across species in the same behavior (i.e., execution). We found that the different behavioral demands of motor preparation increased dynamical distance relative to the across species comparisons during movement execution, even in the same neural population on the same set of trials. The distance became highly significant when comparing across individuals (**Fig. 4e**). Interestingly, many session pairs which were not well-fit by the DSA procedure (**Fig. 4f**, magenta dots), indicating that even if dynamical distance was apparently low, motor preparation activity may not follow the same dynamical rules as execution activity. We did not find dissimilar dynamics when comparing different phases during movement, such as the reaching and pulling phases (**Fig. S6a**), underscoring the altered dynamics are due to changing computations and not the passage of time or differences in measurable behavioral output. Together, these results illustrate that different behavioral demands have a greater impact on neural dynamics for the execution of limb movements than millions of years of evolution.

### Low dynamical distance predicts the generalization of dynamical systems models

Low dynamical distance between pairs of transition matrices **K** indicates that the models represent similar dynamical systems. Intuitively, we expect that models fit to neural activity on one session should better generalize to reconstruct the activity of other sessions if the dynamical distance between them is low. We confirmed this intuition by comparing how well each aligned transition matrices **CK_b_C^-^**^1^ reconstructs the neural activity of the target session in the linearized dynamics space (**Fig. 4f, S7**). We found an orderly, monotonic relationship between this reconstruction performance and dynamical distance, with low dynamical distances yielding the best reconstruction. The models exhibiting the worst generalization were the comparisons across brain regions or across behavioral phases (motor preparation and execution). Thus, low dynamical distances indeed correspond to the good generalization of dynamical systems models fit to the neural population data, confirming that the low dynamical distance across species likely indicates conserved neural computations.

### Varying behavioral output corresponds to different neural geometry

The conservation of neural dynamics across dynamics is particularly surprising given the profound variation in the underlying behavior (differences in grasp types, differences in task design and goals, etc.) to say nothing of the striking differences in limb effectors across these species. Intuitively, motor cortical activity should change in some way to reflect these differences in behavioral output. What, then, accounts for the strong conservation of neural dynamics across the three species? While dynamics capture the underlying rules governing the unfolding of trajectories over time, they are not the complete picture. Trajectories are also described by their geometry, or the shape of their path along the manifold^20,22,37,38^. It is thus possible for the same dynamical system to produce different outputs simply by tracing different paths through an unchanging flow field (**Fig. 5a**). In contrast, a different dynamical system (different flow field) could in principle produce either geometrically similar or dissimilar trajectories. This distinction makes it critical to evaluate both the geometry and dynamics of neural population activity to obtain a deeper intuition for the conserved computations across species. Previous work has demonstrated that individuals of the same species produce geometrically similar population trajectories when producing the same behavior^21^. However, varying behavior led to variations in geometric similarity of motor cortical activity. In light of this prior work, we hypothesized that these specific variations in behavioral output across individuals and species were expressed in the geometry of the neural trajectories, while still subject to the rules imposed by the conserved dynamics previously identified across the three species.

**Figure 5.**
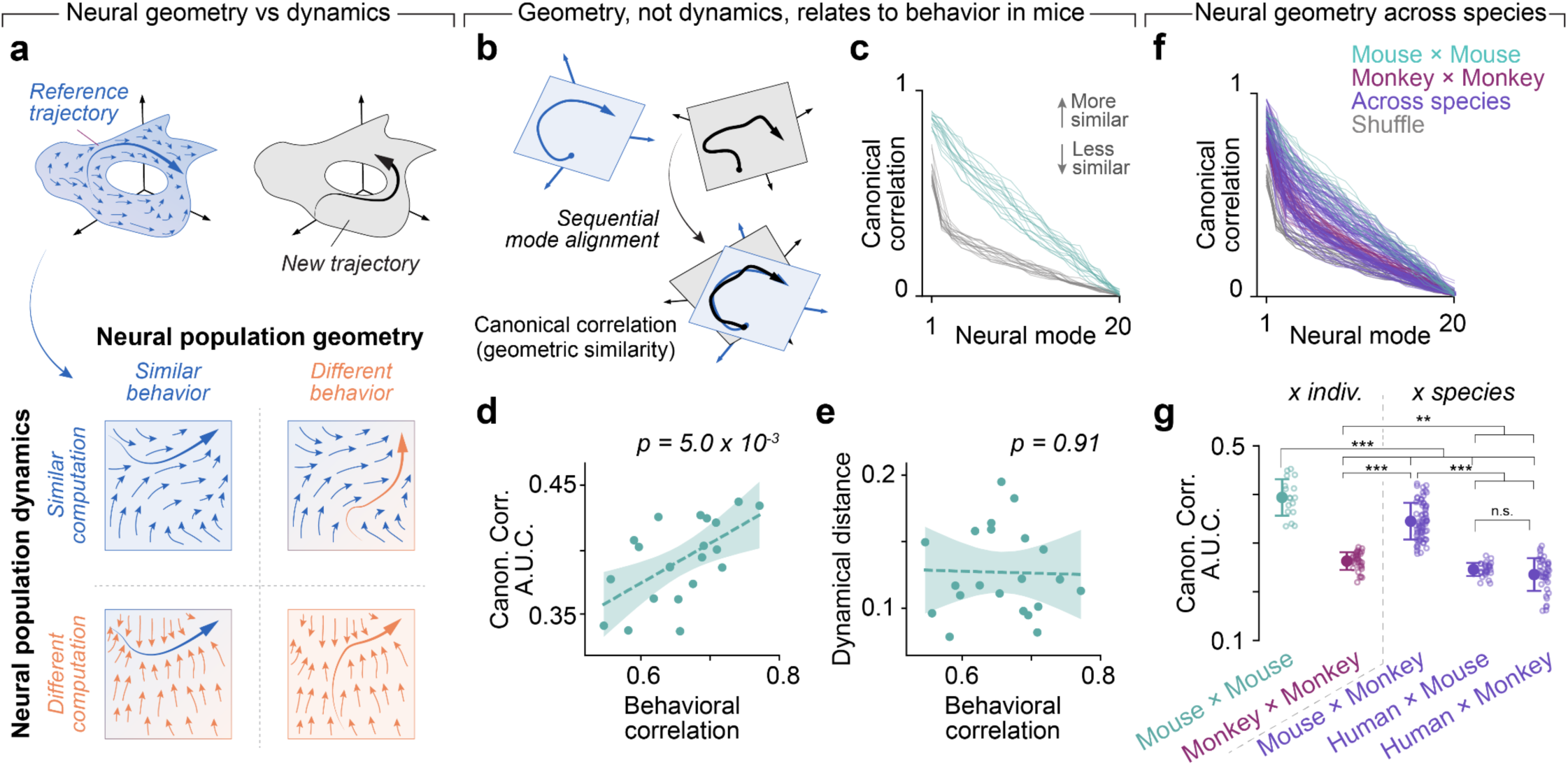
Geometric similarity of neural population activity across species. (a) The similarity of two observed neural trajectories can be described dynamically (the underlying dynamical structure, represented by flow fields) or geometrically (the shape in state space), corresponding to the computations implemented by the population and the behavior specified by the activity, respectively. (b) Canonical correlation analysis (CCA) sequentially aligns new axes (modes) in the neural state spaces of two sets of trajectories, finding a space that maximizes the correlation between the signals. **(c)** Canonical correlation values across mice (green) compared to a shuffle control (gray) that serves as a lower bound (N=17). Due to the sequential alignment, CCA necessarily gives monotonically decreasing correlations for each mode. (d) We computed the area under the curve (AUC) for each set of canonical correlation values for each pair of sessions and plotted these against the average correlation of behavioral covariates (kinematics) for those sessions. Thus, each dot represents a pair of sessions from different mice. Line and shaded error represent a linear regression to the points with 95% confidence intervals on the slope. The *p* value for the slope parameter is shown above the plot. (e) Data presented as in Panel d, but with dynamical distance against the behavioral correlation for each pair of sessions. **(f)** CCA alignment across species (purple) for pairs of monkey and mouse sessions (N=70), human and mouse sessions (N=28), and human and monkey sessions (N=40). Data plotted as in Panel b. (g) AUC for all pairs of sessions across mice (green), monkeys (red), and across species (purple), quantifying the data shown in Panel f. The AUC for the control distributions of each group are also plotted. All statistical comparisons: Wilcoxon’s rank-sum test; *: *p*<0.01; **: *p*<0.001; ***: *p*<0.0001; n.s.: not significant. See Table S1 for exact p-values.

We tested this prediction first in mice, where we had the richest recordings of behavioral (kinematic) covariates, and all individuals were trained to perform the same task. We used Canonical Correlation Analysis (CCA) as a measure of the geometric similarity of neural population activity across sessions, individuals, and species^21,39^ (**Fig. 5b**, **S8**). Consistent with previous work^21^, the geometric similarity was strongly correlated with movement similarity (**Fig. 5c,d**), with more similar behavior (quantified by correlations between kinematic variables, see Methods) corresponding to higher geometric similarity by CCA. Interestingly, we found that dynamical distance was decoupled from behavior (**Fig. 5e**), implying that the different trajectory geometries were implemented by similar dynamical systems.

We next compared geometric similarity across species. Given the different timescales of the human movements compared to those of the mice and monkeys, we removed the effect of time and speed by interpolating the time basis of each trajectory to a fixed length for comparison by CCA. This, in effect, is a best-case-scenario for finding geometric alignment throughout the movement period. We found that across monkey alignment was significantly reduced, likely due to the highly varied behavioral output across the different individuals. Interestingly, monkey to mouse alignment showed some partial similarity due to the structured and repetitive nature of the reach, grasp, and pull tasks that both species performed (and the similar timescales, see **Fig. 2d,e**). In contrast, human trajectories during the object transport task showed almost no alignment to either mice or monkeys, with values near our lower bound control distributions (**Fig. 5f,g**). These results reinforce that the motor cortex can readily produce outputs required for individual- and species-specific behaviors by tuning the path of the trajectories within highly conserved dynamical systems.

### Dynamical similarity arises from shared circuit properties

The above results illustrate two key points: (1) neural trajectory geometry, but not dynamics, directly relates to the similarity of behavioral output; and (2) a conserved dynamical system can still produce varied behavioral output through these geometries. We provided evidence that mice, monkeys, and humans indeed can produce a variety of reaching, grasping, and object manipulation behaviors using the computations of an evolutionarily conserved dynamical system. However, our *post hoc* analysis of neural recordings cannot address two important open questions. First, is it necessarily true that we should see conserved dynamics during the production of reaching movements, or is it plausible that other dynamical systems could produce these behaviors? Second, what is the origin of the conserved dynamics? For this second question, a plausible explanation is that conserved dynamics are the consequence of architectural constraints imposed by anatomy or the neural circuitry shaped by evolution and formed during development.

We designed a simulation—where we have access to the ground truth of circuit parameters—to test how circuit architecture and developmental (or learning) trajectory influence neural dynamics, neural geometry, and behavior. Using the MotorNet toolbox^40^, we trained recurrent neural networks (RNNs) to produce limb movements by driving simulated muscles in a biomechanical model of a primate arm^41^ (**Fig. 6a**). This framework allowed us to fully manipulate architectural variables beyond experimental constraints and study the impact on RNN dynamics and geometry, notably network architecture (e.g., vanilla RNNs vs Gated Recurrent Units (GRUs)^42,43^), learning rate, learning algorithm (e.g., e-prop^44^, feedback alignment^45^), loss function (e.g., different regularization terms), sensory feedback (e.g., manipulations to proprioception and vision), and biomechanical variables (e.g., effector geometry, nonlinear Hill-type muscles vs linear muscle actuators^46^). For a full overview of all explored modifications, see **Table S2**. We simulated each of the 17 configurations 12 times from different random seeds, giving 204 networks to compare. We trained each network to produce random point-to-point reaching movements^47,48^, then instructed them to perform a structured center-out task modeled on typical primate^39,49^ and human^50^ experiments (**Figs. S9**). Compared to the less constrained reaching and grasping tasks analyzed in the previous datasets, this simplified task was advantageous as it provided a clear target in terms of solution optimality based on human psychophysics literature^51^. Further, neural activity during this task is known to give structured, ring-like geometries to neural trajectories^52^, providing a fair benchmark for comparisons when evaluating similarity of neural geometries produced by the networks.

**Figure 6.**
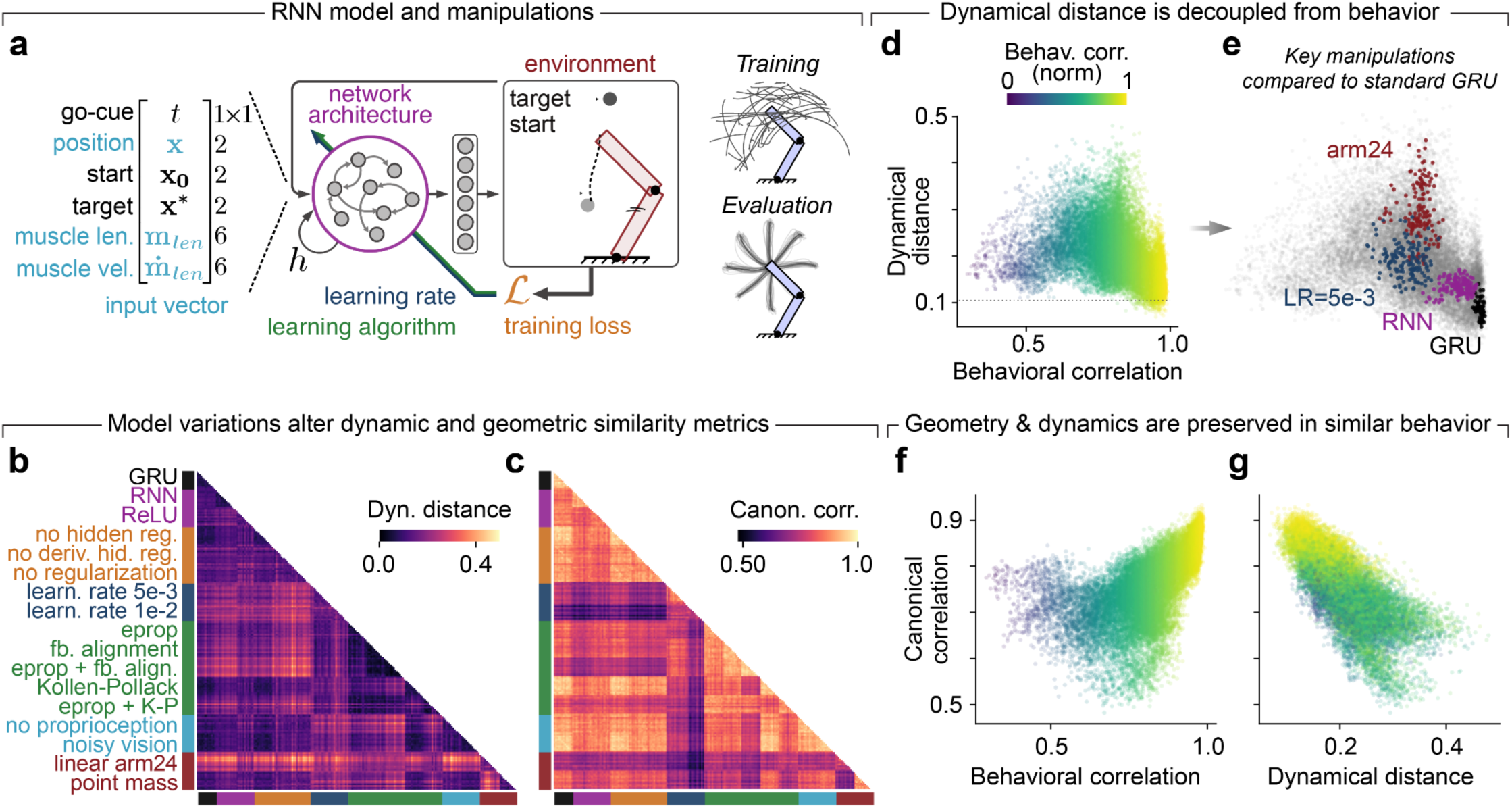
Recurrent neural network models produce a wide range of dynamics and geometries for the same task. **(a)** To study the links between neural circuit architecture and development in a motor control context, we ran biomechanical simulations of a primate arm controlled by a recurrent neural network (RNN). We then manipulated numerous features of the simulation: learning algorithms, loss functions, environment, network architecture, etc (see Methods for details). The models were trained in random point-to-point reaches then evaluated in a center-out task. (b) Heatmap of dynamical distance across all pairs of 12 examples of each of the 17 simulation configurations. Colormap shows dynamical distance where brighter values mean more dissimilar dynamics. Labels are colored by type. Comparisons were made across architectures using the unit activations in the RNN. See Table S2 for full label definitions. (c) Heatmap of canonical correlations for RNN unit activations between all pairs of simulation configuration runs. Colormap shows the area under the curve, with bright colors meaning more geometric similarity. (d) Scatter plot of all datapoints from Panel b against the behavioral correlation (from endpoint velocity) for comparisons to our assumed default configuration, with full biomechanical parameters and GRU networks (see Table S2). Points are colored by the behavioral correlation (colorbar inset). (e) Visualization of specific subsets of architectural manipulations for the scatter plot shown in Panel d: arm24, learning rates of 5e-3, RNNs, and the GRU (the baseline model configuration; see Methods). **(f)** Scatter plot of datapoints from Panel c against behavior correlation. Data presented as in Panel d. (g) Scatter plot of canonical correlations against dynamical distance for all pairs of simulated networks.

We performed DSA and CCA on the RNN unit activations (**Fig. S10**) while producing the center-out movements for all pairs of networks (**Fig. 6b,c**), and compared the similarity in behavioral output using the correlation of endpoint kinematics of the simulated arms. Consistent with our mouse dataset (**Fig. 5e**), there was no significant correlation between dynamical distance and behavioral similarity (**Fig. 6d**). However, we saw substantial variability in the dynamical similarity of solutions produced by the RNNs depending on architectural variations; for brevity we highlight just two of these interesting comparisons. First, simply changing a GRU to a vanilla RNN had a significant impact on both behavioral and dynamical similarity, a demonstration that even seemingly minor modeling decisions could induce qualitative changes in the neural population activity (black and magenta in **Fig. 6e**). Second, we saw large and equivalent impacts on behavioral similarity by removing the Hill-type nonlinearities in the modeled muscles and reducing them to a minimum set of monoarticulars (arm24) and simply changing the learning rate in the cost function (LR) (red and blue in **Fig. 6e**). Yet, these two network architectures produced drastically different dynamics. Notably, the dynamical distance values for different networks with the exact same configuration but different random initializations (black “GRU” dots in **Fig. 6e**)—our best guess for systems which should have similar dynamics for motor control—have similar magnitude and variance as the across species distances in our real motor cortical recordings (**Fig. 3**). The strong dependency of network dynamics on architectural manipulations, and difficulty in producing values observed in our neurophysiological data even in idealized scenarios, reinforces our interpretation that the dynamical distances seen across mice, monkeys, and humans are consistent with conserved dynamics arising from similar circuit properties.

We next studied the interplay between geometry and dynamics as they relate to the behavioral output of the model. Consistent with the mouse datasets, we saw a strong correlation between geometry aligned by CCA and behavioral output (**Fig. 6f**). In other words, the RNNs produced similar behavior through similarly shaped neural trajectories, regardless of architectural configuration and even in the face of varying nonlinearities in the system. We also saw a significant relationship between dynamical distance and geometric similarity in the RNNs, an indication that these two properties cannot be arbitrarily dissociated in our model (**Fig. 6g**). Most intriguingly, the networks that produced the most similar behavior exhibited a high degree of both dynamical *and* geometric similarity. This result underscores the importance of common circuit properties for shared neural computations. Previous work showed a link between behavioral and geometric similarities in both mice and monkeys^21^. Here, we show that achieving both geometric and dynamical similarity during the same behavior can be a challenging task, which is most readily achieved when circuit properties are conserved.

Together, our simulations confirm that: (1) geometry, but not dynamics, must be conserved to produce similar behavior (**Fig. 6d,f**); (2) underlying dynamics need not be similar to produce different behaviors (**Fig. 6d**); and (3) similar architectural elements are necessary to produce both dynamical and geometric similarity during the same behavior (**Fig. 6g**). We thus interpret the low dynamical distances during movement execution in mice, monkeys, and humans to be indicative of conserved neural computations resulting from shared circuit architectural elements^5^, likely inherited throughout evolution in the motor cortex.

## Discussion

We studied mice, monkeys, and a human participant all performing variations of a motor behavior centered on reaching, grasping, and object manipulation. We sought evidence for conserved neural computations in the motor cortex—an evolutionarily old cortical region—across these species. We identified clear evidence for conservation of neural dynamics —our proxy for the computations implemented by a circuit—across species during these motor behaviors. Importantly, we show that such dynamical similarity in the motor cortex across all three species exceeds two critical controls where we can reasonably assume the circuits implement different computations: across two functionally and anatomically distinct brain regions that both critically underlie dexterous movement (motor and somatosensory cortex) in humans, and between motor preparation and movement in monkeys (even when comparing the same neural population). We propose that individual-and species-specific behavioral idiosyncrasies could be readily encoded in the specific geometries of the neural population trajectories, subject to the constraints of the conserved dynamical system that governs motor cortical computations across species.

Our study represents, to our knowledge, the first direct comparison of the dynamics of motor cortical population activity across species, from mice to humans. The dynamical similarities we identify are particularly surprising for two main reasons. First, while there is considerable evidence for cellular and anatomical similarities across species^5,11^, mice and primates have a number of notable differences^5,16–18^ that enable each species’ necessary behavioral adaptations. Our results indicate that these anatomical elaborations enable increasingly complex behaviors while preserving the underlying neural computations. Second, there are a number of notable differences in effector properties (e.g., body mass, limb length, hand and shoulder anatomy) which pose different control problems^53^. How, then, should we interpret the dynamical similarity across species? We believe we have identified an invariance at the level of neural populations which dictates the computations of the motor cortex. Invariance to the finer-grained details of the neural substrate (e.g., particular sets of neurons or circuits) seems to be a key feature of the adaptive processes that govern the behavior of individuals from the same species^21,54^, such as those mediated by the neural computations explored here. While geometric invariances have been previously described across individuals of several species^21,37,38,55^, here we go further and demonstrate invariances in dynamical rules both across individuals and species within the motor cortex.

Despite a growing number of studies exploring the geometry or dynamics of neural population activity independently^20,32,56–59^, the relationship between these properties are rarely directly compared for behavioral control. Our RNN simulations (**Fig. 6g**) demonstrate that neural geometry and dynamics appear well-coupled during relatively simple behavioral tasks such as center-out reaching: networks that produced the most similar behavior had the most geometric and dynamic similarity. However, our across-species comparisons suggest that more complex and finely differentiated behaviors could still be implemented through changes in geometry without needing to alter the underlying dynamics. In this study, we treat geometry only by the shapes of the time-varying trajectories we observe. Recent studies have uncovered structural invariances in neural manifold geometries across individuals and even sleep/wake states^37,38^. Such manifold geometries constrain the possible states that the neural trajectories take, and such structure would be influential on our CCA analyses if it existed in the motor cortex. While our present work does not directly dissect whether geometric structure stems from such fundamental manifold constraints (e.g., activity is confined to live on a specific surface embedded in population activity^20^), our RNN simulations assert that geometry is likely shaped by external inputs. For example, the largest differences in geometry across architectural variations (**Fig. 6e**) stemmed from altered feedback inputs (e.g., arm24) or noisy learning over those inputs (the learning rate manipulations). This points towards a potential mechanism for biological brains to adapt behavior through input-driven reconfiguration of network trajectories without modifying the underlying dynamics, consistent with prior motor learning work^60^.

We focused on the motor cortex due to its evolutionary lineage, ubiquity in mammalian motor control, and direct, observable relationship with motor output. Accordingly, we study three species who all have arm or forelimb effectors with qualitatively similar properties, such as elbow joints that constrain the reaching profiles, and hands that allow for grasping objects. Undoubtedly, there are species specific behaviors enabled by neural expansions or alterations that are not going to be translatable across many species, such as the massive expansion of frontal cortex from monkeys to humans^61^, or the rodent barrel cortex for whisking^62^. Furthermore, while we focus here on voluntary forelimb/arm reaching movements, there is evidence that qualitatively different behaviors involving those same limbs such as quadripedal locomotion may be implemented through different computations^63,64^. A taxonomy across behaviors and species at the level of neural computations, capturing both convergences and divergences, would be a fascinating topic for future study to understand how universal dynamical systems features may be across the phylogenetic tree.

The conservation of neural computations across species has strong implications for translation of animal studies to our understanding of human brain function. Access to neural implementations of these computations at the population level in humans have historically been difficult, necessitating extrapolation from animal models. We believe shared dynamical mechanisms can accelerate both fundamental scientific progress and clinical translation of interventions at the neural population level. Neuromodulation techniques, for example, are often developed in animal models^65^ before deployment in human populations^66^. We foresee that conserved computational implementations across species will enable the development of multi-species neural foundation models^67,68^ with broad implications for neuroscientific and clinical fields such as neural interfaces^69^.

## Acknowledgements

We thank Dr. Jon Sakata and Dr. Raeed Chowdhury for helpful comments on early drafts of this manuscript. A.R.S., Z.C., and N.G.H. would like to thank the Rehab Neural Engineering Labs (RNEL) at the University of Pittsburgh for regulatory support. We thank the following funders for their support of this work. O.C. received funding from the Centre UNIQUE and the Centre Interdisciplinaire de Recherche sur le Cerveau et l’Apprentissage (CIRCA). J.T.D. is a Senior Group Leader at Janelia Research Campus of the Howard Hughes Medical Institute. The human study was supported by the National Institute of Neurological Disorders and Stroke, UH3 NS107714, R01 NS131953 and R01 NS130302. J.A.G. received funding from the European Research Council (Grant, ERC-2020-StG-949660), and the Advanced Research and Invention Agency (Grant, SCNI-PR01-P09). M.G.P. received funding from the chercheurs-boursiers en intelligence artificielle program from the Fonds de recherche du Quebec - Santé, the Collaboration on Motor Planning, Execution, and Resilience (COMPERE), a Natural Sciences and Engineering Research Council of Canada Discovery Grant (RGPIN-2025-06184), and a Canadian Institutes of Health Research Project Grant (PJT-203712).

## Author contributions

Project conception: O.C. and M.G.P. Supervision: G.L. and M.G.P. Neural data analysis: O.C. and M.A. Modeling and simulations: O.C.

Figure preparation: O.C., M.A., and M.G.P. Manuscript writing: O.C., M.A., and M.G.P.

Manuscript editing: O.C., M.A., A.R.S., J.T.D, J.A.G., G.L., and M.G.P.

Data collection: A.R.S., Z.C., J.P., N.G.H, and J.T.D.

## Data & code availability statement

The mouse dataset is publicly available from Janelia Research Campus (https://doi.org/10.25378/janelia.28025282.v1). The two monkey datasets are publicly available, the first on Dandi (Monkey 1: https://dandiarchive.org/dandiset/000941; Monkey 2: https://dandiarchive.org/dandiset/001209) and the second on GIN (https://doi.org/10.12751/g-node.f83565). The human dataset will be made available on DABI upon publication. Code to reproduce the analyses and simulations for this paper will be made available when ready or upon publication.

## Methods

### Mouse behavioral task and neural recordings

We analyzed publicly available behavioral and neural recordings of mice performing a reaching and pulling lever task. For detailed experimental methods on the experimental setup and data processing, see Ref. 24. In brief, four 8-16 week old mice were trained to reach for and pull on a lever over a period of approximately one month before undergoing surgical procedures to open a craniotomy. For the analyses of this manuscript, we analyzed seven total sessions. During recordings, mice self-initiated a reaching movement after the lever appeared. The lever was placed 1.5 cm away from the resting hand position. The task involved four trial types: the lever could be placed in one of two positions (left or right) spaced approximately 1 cm apart and was loaded with one of two weights (light: 3g, heavy: 6g). The mice completed 20 trials of each trial type in blocks, with trial types switched sequentially, and the cycle was repeated twice to give 160 trials per recording session. Movement kinematics were tracked with markerless pose estimation from video cameras, and neural activity was recorded using Neuropixels probes inserted through a craniotomy targeting the motor cortex. Neural activity was spike sorted with Kilosort 2.0 (https://github.com/MouseLand/Kilosort) and manually curated to identify well-isolated individual neurons. The average number of motor cortical units included for each mouse was: mouse 38, 117± 28 (range, 95–157); mouse 39, 64; mouse 40, 77 ± 4 (range, 73–81); and mouse 44, 55. These neural spike trains were aligned to the reach onset (extracted from the kinematic tracking) for subsequent analysis.

### Monkey behavioral tasks and neural recordings

We analyzed neural data from two different publicly available datasets of monkeys performing motor tasks. The first dataset (which we refer to as the *cylindrical object dataset*) comprised 15 sessions of recordings from two rhesus macaque monkeys (*macaca mulatta*, Monkey X: 8 sessions and Monkey L: 7 sessions)^28^. For detailed methodology, see the original manuscript in Ref. 25. In brief, the monkeys were trained to reach for, grasp, and manipulate objects (following a delay period and a go cue) to receive a liquid reward. In the full recordings, the monkeys reached from a central object for four different types of peripheral objects arranged on a rotating board. Only two of these objects (the cylindrical objects, but not the sphere and the button) required the monkey to pull on the object to receive a reward. We thus subsampled the trials (‘coaxial trials’) in the full dataset to focus only on this object type for most direct comparison to the mouse task described above, yielding 4 sessions per monkey for subsequent analysis. The rotating board could take 8 different orientations, giving us 8 different trial types for the cylindrical object pull task. As the monkeys performed this task, neural spiking activity was recorded using six chronically-implanted floating microelectrode arrays (16 electrodes per array) implanted in the primary motor cortex along the central sulcus, yet, the available dataset focuses only on the 4 middle one (details on array position can be found in Ref. 70). Spike waveforms were detected from threshold crossings then manually sorted into individual units. We binned these spike trains at 20 ms resolution for subsequent analysis.

The second dataset (which we refer to as the *cubic object dataset*) comprised 2 sessions of recordings from two rhesus macaque monkeys (*macaca mulatta*, Monkey N and Monkey L)^29^. For detailed methodology, see the original manuscript in Ref. 26. In brief, the monkeys were trained to reach for a cubically shaped object and grasp it using either a precision grip or a side grip following a visual cue then pull on it and hold it for 500 ms within a certain position window before receiving a liquid reward. As in the cylindrical object dataset, the monkeys had a delay period before the go cue. The object was loaded with two different loads (low force: 1N; high force: 2N), giving four different trial types. As the monkeys performed this task, neural spiking activity was recorded using chronically implanted Utah microelectrode arrays targeting the arm region of the motor cortex^31^. Spike waveforms were detected from threshold crossings then manually sorted into individual units. We binned these spike trains at 20 ms resolution for subsequent analysis.

For each dataset, we pooled all trial types to give as much behavioral richness as possible when comparing the neural dynamics. For both tasks, we took 600 ms of time before the go cue as a motor planning period for the analyses of Fig. 4c, and we defined the movement period to range from the movement onset (defined as when the monkey lifted the hand from rest) to the movement offset (determined as the reward time in the cylindrical object dataset and the start of the holding period in the cubic objects dataset). We separated the reaching and pulling phases by the grasp time, estimated by either object contact or onset of pulling force.

### Human behavioral task and neural recordings

We analyzed neural recordings from human sensorimotor cortex. The study that produced this dataset was conducted under an investigational device exemption from the US Food and Drug Administration and approved by the institutional review boards at the Universities of Pittsburgh and Chicago. The clinical trial is registered at ClinicalTrials.gov (NCT01894802). Informed consent was obtained prior to any study procedures. The participant (male) was 57 years old at the time of implant and presented with a C4-level American Spinal Injury Association (ASIA) D SCI occurring 35 years before the implant. The study inclusion criteria specified that the participant reported no functional use of their hands and had less than grade 2 strength in finger flexor and abduction muscles. At the time of these experiments, the participant was able to pick up objects using a compensatory tenodesis grasp—voluntary wrist extension that passively pressed digits and the object into the palm, with minimal active contraction of extrinsic or intrinsic finger flexors. Movement times were slower than would be expected for an able-bodied individual. Four Blackrock NeuroPort Electrode (Blackrock Neurotech) microelectrode arrays, analogous to Utah microelectrode arrays, were implanted in the sensorimotor cortex, two in the primary motor cortex (96 channels each) and two in the somatosensory cortex (32 channels each). In the motor cortex, one array targeted the arm region and one targeted the hand region. Both sensory arrays targeted hand regions. We focused our analysis on the more active arrays—lateral motor and medial somatosensory—to ensure the best estimates of population dynamics.

The participant completed 4 recording sessions of an object movement task. Each trial proceeded as follows. A cylindrical object was placed in one of 3 positions on a table. A light ring was illuminated on the table to cue a target location towards which to move the object. One or more obstructions were placed between the initial position and the target location to force the participant to fully lift the object during the movement. The participant began at rest until a go cue signified the trial start. The participant then reached for and acquired the cylinder and transported the object to the target. The cylinder was instrumented with force-sensitive resistor sensors on 4 sides to track the object acquisition time, and a force sensor on the bottom to track the liftoff. The data from the sensors was synchronized with the simultaneously acquired neural data. The weight of the object was changed on each trial by adding or removing metal weights inside the cylinder.

As the participant completed the behavioral task, we recorded intracortical activity from the implanted Utah arrays. We extracted multi-unit spiking activity from threshold crossings on each of the motor and somatosensory cortical microelectrode arrays. The threshold was defined as -4.5 standard deviations from the mean. Previous work has demonstrated that sorted single units (such as those in the mouse and monkey datasets) and multi-unit threshold crossings provide equivalent estimates of neural latent dynamics^71^, enabling direct comparison across these datasets. We aligned the neural data to the go cue for subsequent analysis. However, since the trial duration was significantly longer in the humans compared to the mice and monkey datasets, we devised a strategy to afford more comparable movement windows. We isolated a period throughout the trial based on the object acquisition time, taking 0.8 s before and 2.4 s after. (see **Fig. S1**). As a control, we compared 1.6s windows taken from throughout the trial surrounding object hold, some biased more towards the period before object hold and some biased more towards object transport (see **Fig. S6d,e**).

### Behavioral quantification

For all datasets from all three species, we split the behavioral task into two movement phases: a reaching phase and an object manipulation phase (level pull for mice, object pull for monkeys, and object movement for humans). For each trial, we computed the time spent in each phase and pooled these values across all sessions from each individual for the distributions of **Figs. 2**, **S1** The mouse dataset contained detailed, continuous forelimb kinematics extracted from video recordings. We computed behavioral similarity for the comparisons in Fig. 5d**,e** across trials using correlations in the hand trajectory kinematics (Pearons’s *r*) across pairs of sessions in the mouse dataset.

### Processing of neural spiking data

For each neuron recorded on a given trial, we turned the spike train into a smooth firing rate by applying a square root transform (to stabilize variance) and convolving that spike train with a Gaussian kernel of width 50 ms^72^. To visualize single trial population activity (e.g. **Figs. 2, S2**), we “normalized” each neuron’s firing rates by subtracting the calculated average across trials, subtracting each trial by the trial’s minimal value, and finally scaling the result by the maximal value found over trials. These were visualized as a heatmap over time for all recorded neurons on that trial. To specifically target the movement epoch, we restricted the neural activity between the reaching onset and pulling offset, then concatenated across all trials on a given session and used Principal Components Analysis to extract neural trajectories (e.g. **Figs. 2, S2**). For all primary analyses in this manuscript, we opted for a 20-dimensional latent space for dynamics comparisons which was sufficient to explain 50-100 % of variance explained neural variance across datasets (**Fig. S3**), though our results were qualitatively unchanged for reasonable ranges of assumed dimensionalities (**Fig. S6c**).

### Dynamical similarity analysis (DSA) to compare neural dynamics

#### Fitting dynamical systems models to neural data

DSA is designed to compare the dynamical similarity of two systems such as neural populations. For detailed descriptions of the theory and validation of the method, see the original study in Ref. 23. In brief, DSA involves computing a finite-dimensional approximation of the Koopman operator via dynamic mode decomposition (DMD)^73,74^ on time-delayed embeddings of the neural data. This Koopman operator corresponds to a (usually higher-dimensional) linearization of the transition function between states of a system, here the neural state, allowing us to compare differences between state transition properties of different systems with linear similarity methods^23^. There are three key hyperparameters to consider when performing DSA for neural populations: the assumed dimensionality of the neural trajectories (in our case found by PCA) before performing DSA, the number of delays in the time delay embedding for linearization, and the rank of the DMD. For primary analyses in this manuscript, we set these hyperparameters as 20, 10, and 75, respectively. Importantly, our conclusions were qualitatively unchanged with reasonable variations in these hyperparameters (**Fig. S4**). We selected these values empirically through a cross-validation procedure on the reconstruction performance of the DMD on the neural data. We varied the model’s number of delays from 1 to 14, and rank from 10 to 150, and tested each parameter combination on three assumed dimensionality of the neural trajectories: 15, 20, and 25. We prevented the combination where the rank exceeds the number of delays time the lag, as the model is suggested to overfit the data in such cases^23^. We trained the model on 9 of the 10 folds (constituting the training set) and evaluated the model’s capacity to accurately predict the remaining test fold (ground truth) by computing the correlation (Pearson’s *r*) between the original dataset and the reconstruction from the aligned **K_b_**. By construction, the prediction array integrates the first time points of the ground truth to compensate for the delays, hence, we computed the statistics from the point in time where the model’s prediction starts. We repeated the process of training-testing-evaluating to test predictions on each of the 10 folds.

#### Quantifying dynamical similarity

We computed the dynamical distance (a proxy for dissimilarity of the dynamical systems) on the Koopman operators (**K_a_** and **K_b_**) as a Procrustes dissimilarity over vector fields^23^. In brief, the two state transition matrices are aligned by learning a matrix **C** which linearly transforms **K_b_** to minimize the distance from **K_a_** subject to **CK_b_C^-^**^1^. Note that the **C**s matrices are most likely not symmetric in that sense, we considered a **C** matrix and **K_b_** to reconstruct **K_a_** and another **C** matrix and **K_a_** to reconstruct **K_b_**. Hence, each direction of comparison for a pair of sessions has a unique alignment matrix **C**. We report only one direction to avoid duplicating session pairs in our statistical comparisons, but we saw no qualitative change in results across the board if we reversed the direction (**Fig. S6b**) or included both directions (data not shown). For **Figs. 3d,e**, we used UMAP^33^ to embed the aligned **K_b_** matrices across all datasets for visual comparison. More quantitatively, the dynamical distance is the Froebenius norm of the difference after minimization, where larger values correspond to more dissimilar dynamics. The values are arbitrary and require contextualization. To provide an upper bound on these values, we implemented a shuffle control which was intended to give no correspondence between dynamics. We randomly selected windows of data from each section of the recordings unaligned to each other and to any task variables, ensuring we maintained the same overall statistics of neural activity. To provide a lower bound, we compared dynamics across sessions in the same individual, where we can be reasonably certain the same dynamical system generated the neural data.

### Canonical correlation analysis (CCA) to compare neural geometries

We quantified the geometric similarity in neural trajectories across animals using CCA between pairs of datasets^21,39^. CCA requires point-by-point alignment in time between the two trajectories. Given the large differences in trial time between the human and animal datasets (**Figs. 2**, **S1**), we thus chose to interpolate the trajectories onto a uniform time basis of 100 time points. Our comparison thus provides a “best case scenario” on the similarity of neural trajectories, as we have removed time as a factor and we only require that the trajectories have similar geometric organization. As a lower bound on the correlation values, we computed a null distribution by an identical shuffling procedure to that of the DSA analysis above. For Fig. 5g, we summarized the canonical correlation values by computing the area under the curve (AUC) for each trace.

### RNN simulations to explore architectural influence on geometry and dynamics

#### Architecture

All neural network models were implemented using the open-source Python package pytorch^75^. Models consisted of a recurrent neural network (RNN) composed of gated recurrent units (GRUs) with *tanh* non-linearity or vanilla RNNs. The GRUs were implemented as:

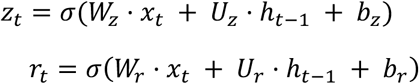

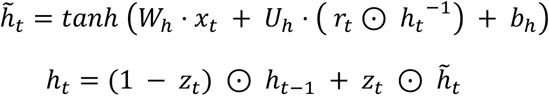

The symbol ⊙ indicates the element-wise product, ℎ^/^_𝑡_ the candidate hidden state at timestep 𝑡, ℎ_𝑡_ the hidden state at timestep

𝑡, 𝜎 the sigmoid function, 𝑈_𝑧_, 𝑈_𝑟_, 𝑈_ℎ_, 𝑊_𝑧_, 𝑊_𝑟_, 𝑊_ℎ_ are learnable weight parameters, and 𝑏_𝑧_, 𝑏_𝑟_, 𝑏_ℎ_ are learnable bias parameters. For the network with a 𝑅𝑒𝐿𝑈 activation function, the 𝑅𝑒𝐿𝑈 function replaced the 𝑡𝑎𝑛ℎ function in the above. The vanilla RNNs were implemented as:

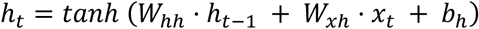

𝑊_ℎℎ_, 𝑊_𝑥ℎ_ were learnable weight parameters and 𝑏_ℎ_ was a learnable bias parameter.

The RNN layer then projected into a fully connected output layer with as many neurons as actuators in the effector and a sigmoid non-linearity to ensure positive outputs. Each output from this last layer served as an action to be passed to one of the actuators controlling the effector. The RNNs were composed of sixty-four units. They received as input a vector containing a step go cue signal, the cartesian position of the effector’s endpoint, the start position and target position, as well as each actuator’s length and velocity^40^ (Fig. 6a). Variations included models where actuator length and velocity were omitted (no proprioception models), and where a Gaussian noise process (𝜎 = 1) was added to the cartesian position of the effector’s endpoint (noisy vision).

#### Environments

The environment models employed were from the open-source Python package MotorNet^40^. The simulation time constant was set to 10 ms for all models. For the control model, we employed the *RigidTendonArm26* effector with default parameter values and the *Runge-Kutta 4* algorithm for numerical integration. The actuators were *MujocoHillMuscle* object with no passive force contribution and 𝑙_𝑚𝑖𝑛_ = 0.3, 𝑙_𝑚𝑖𝑛_ = 1.8.

For the point mass simulations, we employed the *PointMass24* skeleton and set its mass to be equal to the combined mass of both arm segments (1.82 + 1.43 = 3.25 𝑘𝑔). The point mass was actuated by a *ReluMuscle* object with time constants set to the same values as the *MujocoHillMuscle* (𝜏_𝑎𝑐𝑡_ = 0.01, 𝜏_𝑑𝑒𝑎𝑐𝑡_ = 0.04) and maximal isometric force of 1000 N. The point mass actuators each acted on one of the four possible directions (up, down, left, right), with no anchor point and a moment arm of 1 regardless of the point mass position. The point mass was constrained within a 60 cm distance to the origin for all coordinates, forming a square of 1.2 m centered on the (0, 0) origin. Because muscle length and velocity are not relevant for the *ReluMuscle* object, these inputs were omitted for the point mass simulations.

For the “arm24” simulations, we employed the same two-link arm skeleton—specifically, a *TwoDofArm* skeleton object— as for the *RigidTendonArm26* simulations and a *ReluMuscle* actuator with the same parameters as for the point mass simulations. Unlike the point mass, the four actuators acted on joints directly instead of on cartesian positions, with an agonist and antagonist for the shoulder and elbow joint each.

#### Training procedure

All models were trained to reach from a random start position to a random target of 1 cm radius drawn uniformly within the full joint space. Actuator endpoints were required to stay inside the start position until the go cue signal, and to go to the target afterwards. During training, the go cue signal was drawn from a uniform distribution bounded between 0 and the full duration of the episode. Episodes lasted one second, corresponding to 100 timesteps of 0.01 seconds. The instantaneous loss (loss at timestep 𝑡) was:

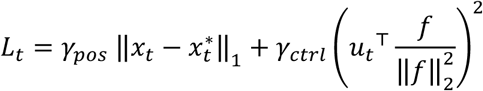

with 𝛾_𝑝𝑜𝑠_ = *0* and 𝛾_𝑐𝑡𝑟𝑙_ = *1* if ‖𝑥_𝑡_ − 𝑥*_t_*^∗^‖*_2_* < 𝑟, 𝛾_𝑝𝑜𝑠_ = *1* and 𝛾_𝑐𝑡𝑟𝑙_ = *0*.*05* otherwise. 𝑥_𝑡_, 𝑥*_t_*^∗^, 𝑢_𝑡_,𝑓 were the position, target, action, and maximum isometric contraction vectors used for normalization of the control penalty, respectively. Therefore, the theoretical lower bound of 𝐿_𝑡_ was 0. All models were trained for a minimum of 20,480 training steps, and then until convergence, defined as all of the last 20 training steps having an average loss 𝐿_𝑡_ under 0.04.

#### Learning algorithms

All models used a learning rate of 3e-4, gradient norm clipping of 0.5, and a batch size of 32. We simulated a full episode for each batch before performing a backward pass and network weight update. We also added a L2 regularization on neural activity and its derivative to the gradient for both model groups to promote parsimonious and sparse network activation levels, which usually leads to more biologically plausible activity patterns^76^. Regularization weights were 𝛾*_ℎ_* = *0*.*01* and 𝛾*^ℎ^* = *0*.*1* for neural activity ℎ and its derivative ℎ, respectively, such that:

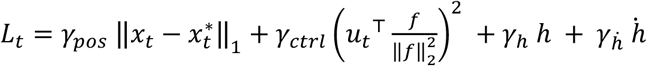

Models were initialized with Xavier uniform and orthogonal initialization with a gain of 1 for input and recurrent weight matrices, respectively. The output layer’s weight matrix was initialized with a Xavier uniform initialization and a gain of 1. Seeds (n=12 per group) were matched between groups to ensure any difference between models did not arise from differences in initial learnable parameter values. Biases were all initialized at 0 except for the output layer, whose bias was initialized at -5. This negative bias ensures that on initial training epochs, output activity is falling well into the low saturating part of the sigmoid non-linearity, leading to action vectors 𝑢_𝑡_ with a lower norm. In practice this greatly helps learning by promoting a stable initial regime of activity^40,77^.

For a comprehensive list of all model variations explored in this study, see **Table S2**.

### Statistical tests

All statistical tests were two-sided, nonparametric tests (Wilcoxon’s rank-sum). We set three stringent tiers of significance on the resulting *p* value: *: 0.01, **: 0.001, and ***: 0.0001. All comparisons with *p*-values greater than 0.01 were deemed not significant (n.s.).

---

**Figure S1.**
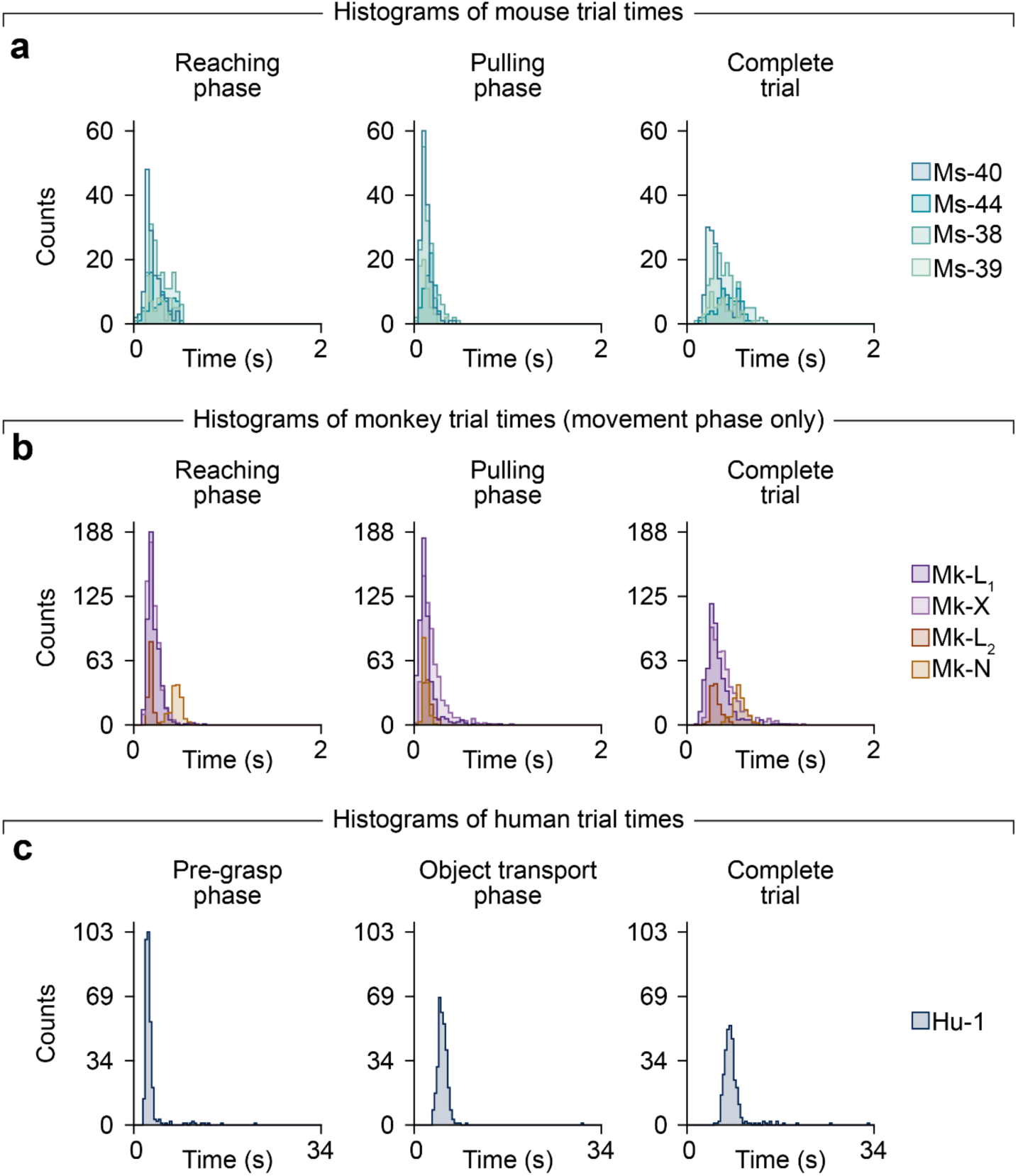
Detailed quantification of trial times across species-specific behavioral tasks. (a) Histograms of the duration of the reaching phase (left), object interaction phase (middle), and full trial (right) for all trials, pooled across all sessions for each mouse (3 for Ms-38, 2 for Ms-40, 1 each for Ms-39 and Ms-44). (b) Trial time histograms pooled across all sessions for each monkey (4 for Mk-L_1_, 4 for Mk-X, 1 each for Mk-L_2_ and Mk-N). (c) Trial time histograms pooled across all sessions for the human participant (4 for Hu-C1). Due to task differences and the partial spinal cord injury, movement phases were significantly longer in humans than the other species. To compare neural population features on similar timescale, we isolated portions of the movement activity surrounding the time of object acquisition (see Methods).

**Figure S2.**
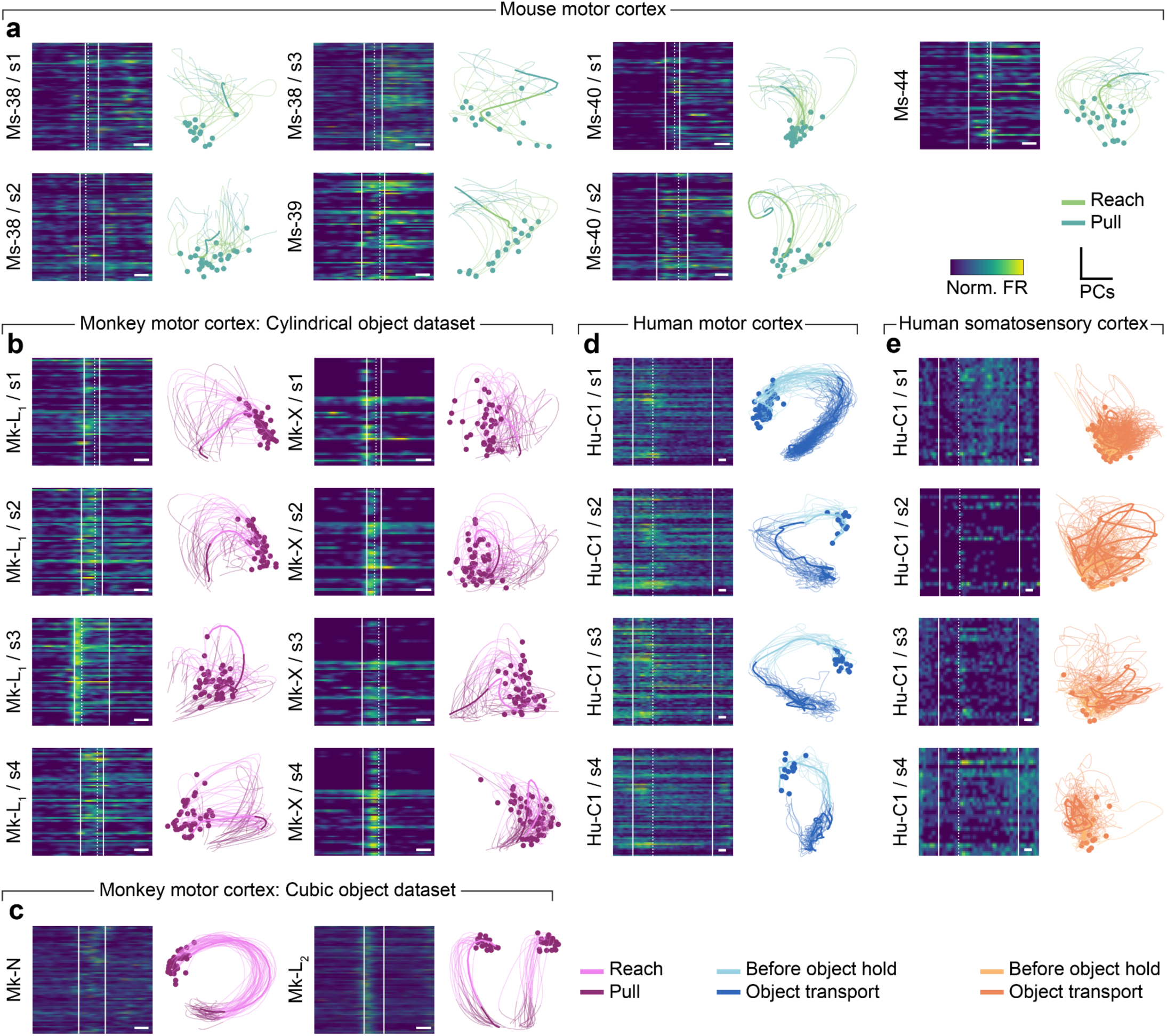
Example neural activity from all mouse, monkey, and human datasets. **(a)** Left: heatmap of neural activity for a single representative trial. Solid lines indicate the start and end of the movement analysis window, and dashed line indicate the time of object grasp. White scale bar indicates 200 ms. Right: example neural trajectories (right) for randomly picked trials (⅓ of total from the session) for all mice sessions. The thicker line indicates the example trials shown in the left panel. (b) Example neural activity for all monkey motor cortical sessions in the cylindrical object dataset. (c) Example neural activity for all monkey motor cortical sessions in the cubic object dataset. (d) Example neural activity for all human motor cortical sessions. (e) Example neural activity for all human somatosensory cortical sessions.

**Figure S3.**
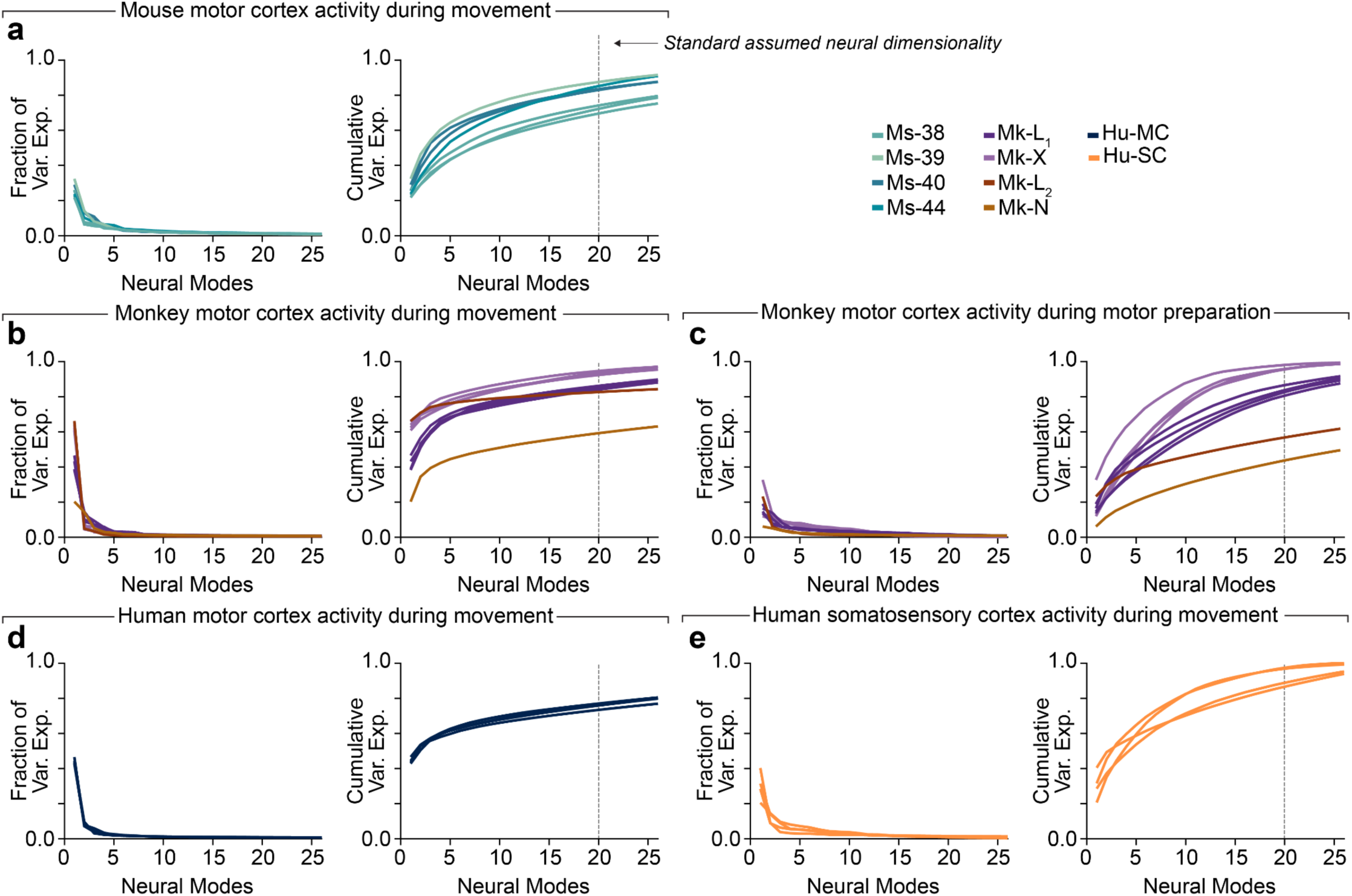
Variance explained by PCA dimensions for all mouse, monkey, and human sessions. (a) Fraction of variance explained (left) and cumulative variance explained (right) by each of the neural modes identified by PCA for all the mouse sessions recorded in the motor cortex during reaching and pulling. The gray vertical line in the cumulative plot indicates 20 PCs, our standard dimensionality chosen for subsequent analyses. (b) Neural variance explained for the motor cortex for all monkey sessions during the movement phase (c) Neural variance explained for the motor cortex for all monkey sessions during the preparation phase. (d) Neural variance explained for the motor cortex for all human sessions during the movement analysis window. (e) Neural variance explained for the somatosensory cortex for all human sessions during the same window as Panel d.

**Figure S4.**
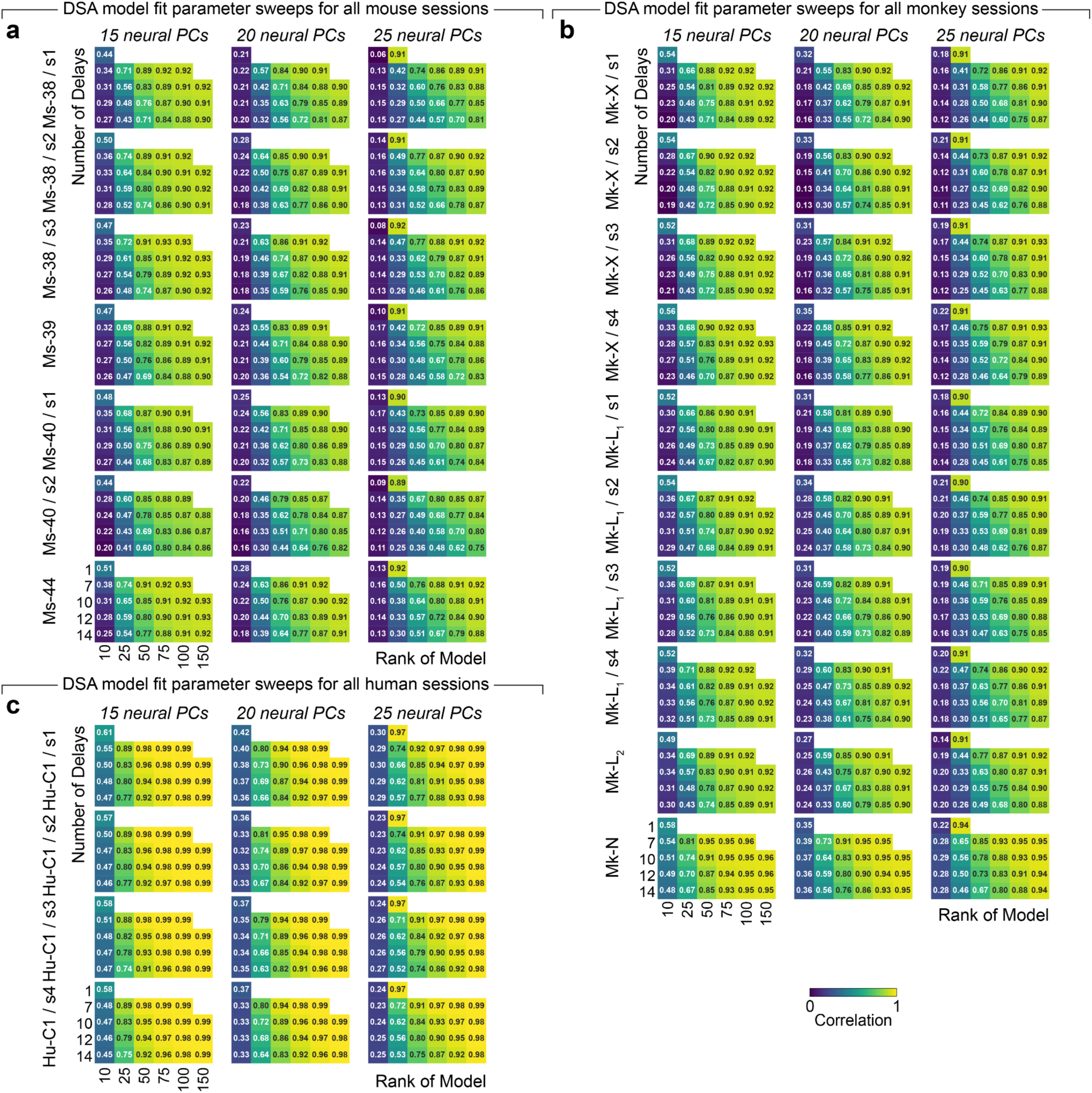
Parameter optimization for DSA models across all datasets. Heatmap of reconstruction performance (correlation) of DSA model fit evaluated for every out of the (a) mouse sessions, (b) monkey sessions, and (c) human sessions for 3 assumed dimensionality (15, 20 and 25 PCs) of the neural space. Heatmap grids show combinations of the rank of the model and the number of delays.

**Figure S5.**
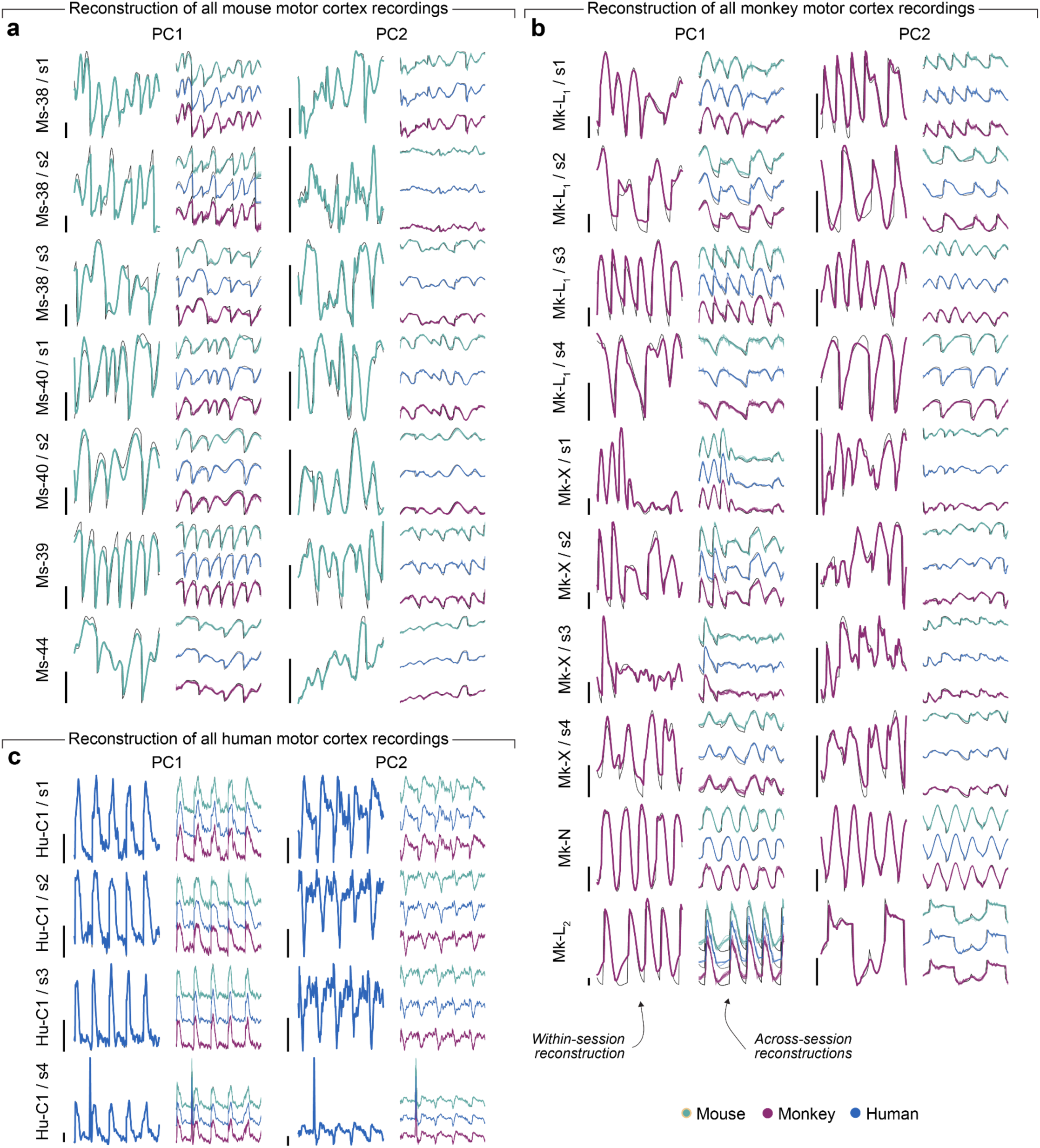
Reconstruction performance across species using DSA model fits. Example traces of the first 2 Principal Components (PCs) for neural data (black lines) for 5 example trials and reconstructed from dynamical systems fits from **K** (colored lines, green: mouse, red: monkey, blue: human) for all mouse **(a)**, monkey **(b)**, and human **(c)** sessions. For each session, we show the reconstruction from the **K_a_** fit on that session (left), and from the aligned **K_b_**s (right) separated by species. Traces are not proportionally scaled to enable more detailed views of the lower-variance components. Black scale bar to the left of the **K_a_** reconstruction shows arbitrary units of 50 within the PC space.

**Figure S6.**
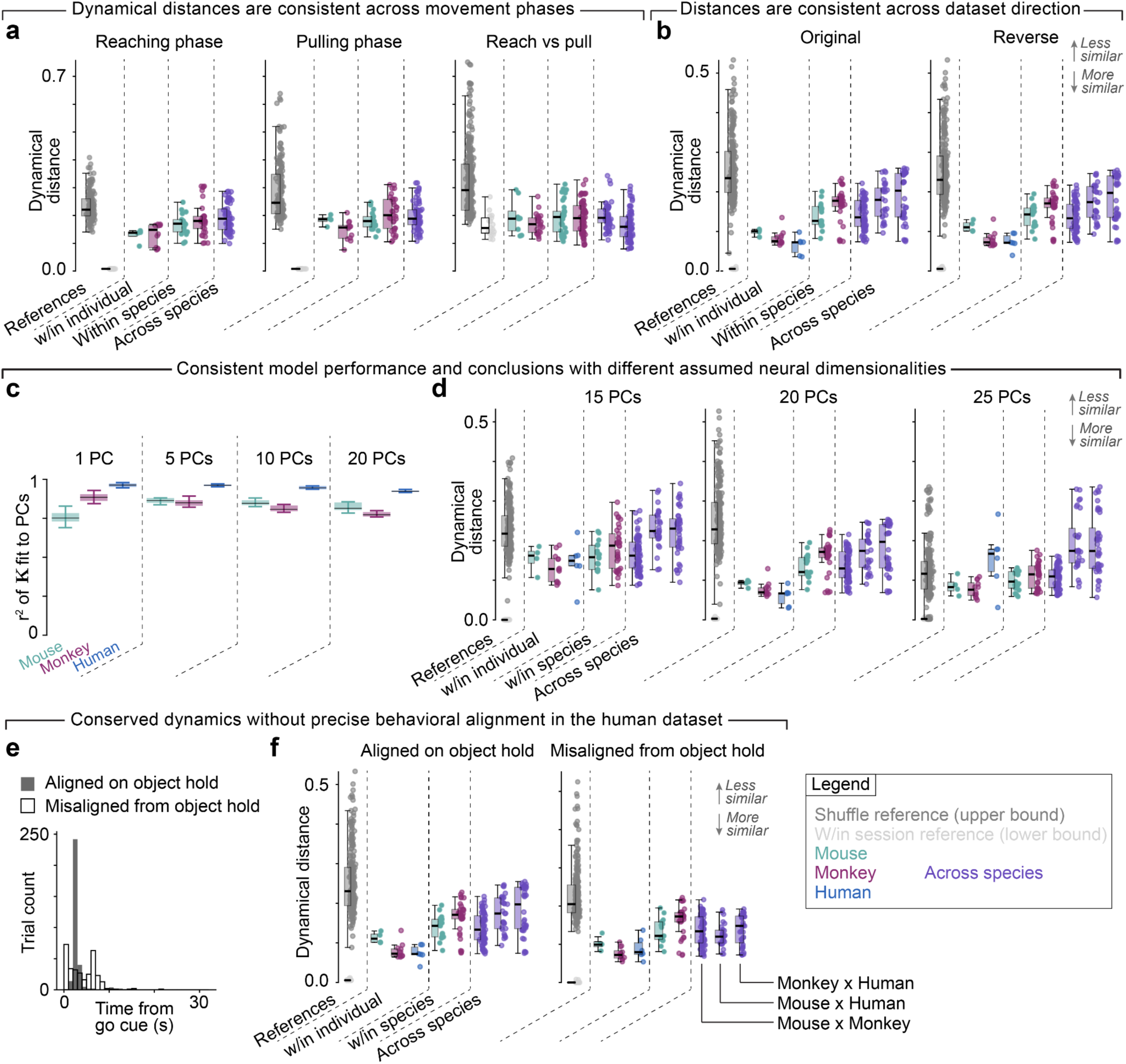
Control analyses for dynamical similarity across species. (a) We ensured dynamical distance values held when separating the trials into reaching (left) and grasping (right) phases. Note that given the differences in task design and motor performance we could not cleanly identify reaching versus pulling phases in the human trials. (b) Since dynamical distances from DSA need not be symmetric, we recomputed the results of Fig. 3f reversing the identity of K_a_ and K_b_. We obtained nearly identical results by this procedure, confirming that we get equivalent similarity regardless of which dataset is chosen as the aligned trajectories or the target trajectories in the optimization. (c) Reconstruction R^2^ of K matrices fit to each session of mice, monkeys, and humans for different assumed neural dimensionalities. We were able to effectively model the dynamics at different dimensionalities except in the extremes (e.g., one-dimensional PC trajectories in mice). (d) Dynamical distance values found by DSA on three distinct assumed neural dimensionality: 15 PCs results (left panel), 20 PCs (middle panel; note that this reproduces the results from Fig. 3f), 25 PCs (right panel). (e) Since we isolated only a portion of the total trial in the human dataset, we ensured our results did not depend on this choice by comparing the original windows to those taken throughout the trial misaligned to the time of object acquisition. (f) Dynamical distance results for the original windows aligned on object hold (left) windows and the misaligned (right).

**Figure S7.**
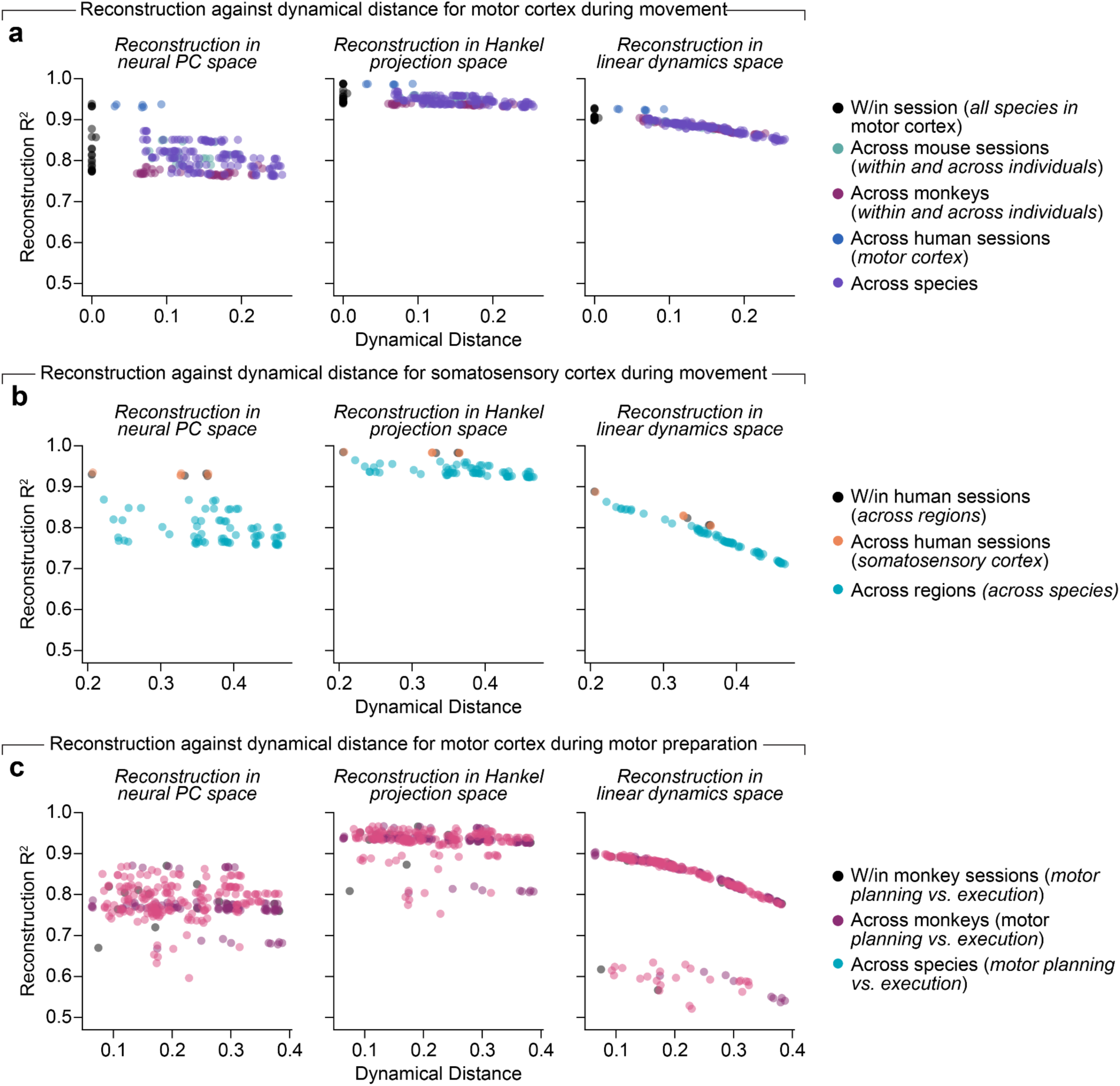
Relationship between DSA model fit reconstruction and dynamical distance. (a) We reconstructed the neural data and its projections at each step of the algorithm: the original neural data in a 20-D PC space (left), the Hankel projection space (middle), and the low-rank linearized dynamics space (right). We show the R^2^ reconstruction performance against the dynamical distance from DSA in the three spaces. (b) Reconstruction against dynamical distances for all pairs of sessions when comparing to the human somatosensory cortex. (c) Reconstruction against dynamical distance for all pairs of sessions compared to the monkey motor cortex during motor preparation. In all plots, note the strong negative relationship in the linearized dynamics space (right panels) between the reconstruction performance and the estimated dynamical distance.

**Figure S8.**
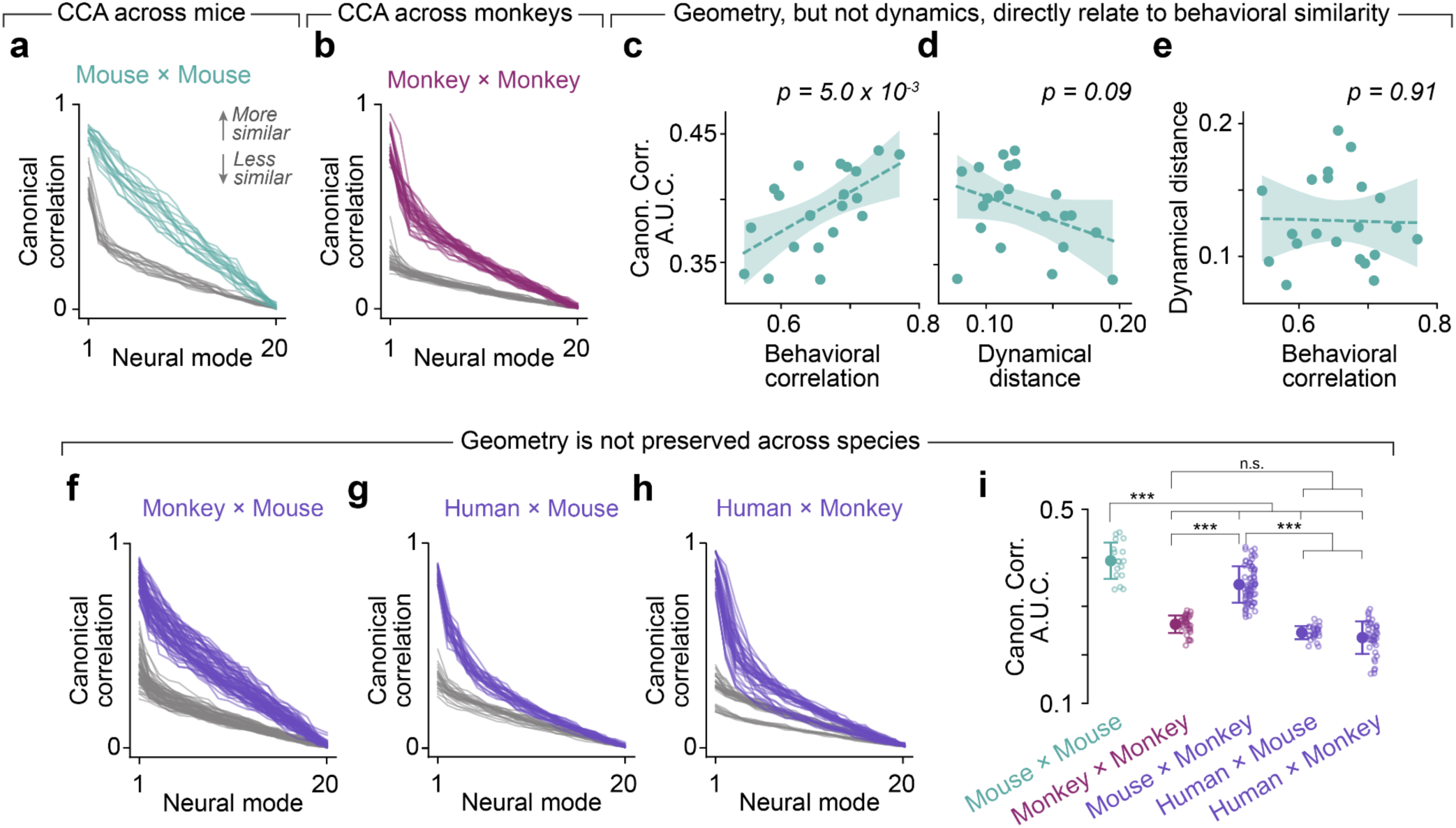
Geometric similarity of neural population activity across species. **(a)** Canonical correlation values across mice (green) compared to a shuffle control (gray) that serves as a lower bound (N=17). Due to the sequential alignment, CCA necessarily gives monotonically decreasing correlations for each mode. **(b)** CCA alignment between pairs of sessions from different monkeys (N=33). Data plotted as in Panel b. Note the noticeable decrease compared to the across mouse results. This outcome is consistent with the close relationship between geometry and behavior, given that our monkey datasets represent a higher amount of behavioral variability (variations in objects and task design, recordings in different lab settings, etc. compared to the mouse dataset, which featured one stereotyped task. (c) We computed the area under the curve (AUC) for each set of canonical correlation values for each pair of sessions and plotted these against the average correlation of behavioral covariates (kinematics) for those sessions. Thus, each dot represents a pair of sessions from different mice. Line and shaded error represent a linear regression to the points with 95% confidence intervals on the slope. The *p* value for the slope parameter is shown above the plot. (d) Data presented as in Panel c, but showing canonical correlation AUC against dynamical distance found by DSA for that pair of sessions. (e) Data presented as in Panel d, but showing dynamical distance against the behavioral correlation for each pair of sessions. **(f)** CCA alignment between pairs of monkey and mouse sessions (N=28). Data plotted as in Panel b. **(g)** CCA alignment between pairs of human and mouse sessions (N=40). Data plotted as in Panel b. **(h)** CCA alignment between pairs of human and monkey sessions (N=70). Data plotted as in Panel b. (i) AUC for all pairs of sessions across mice (green), monkeys (red), and species (purple). The AUC for the control distributions of each group are also plotted. All statistical comparisons: Wilcoxon’s rank-sum test; *: *p*<0.01; **: *p*<0.001; ***: *p*<0.0001; n.s.: not significant.

**Figure S9.**
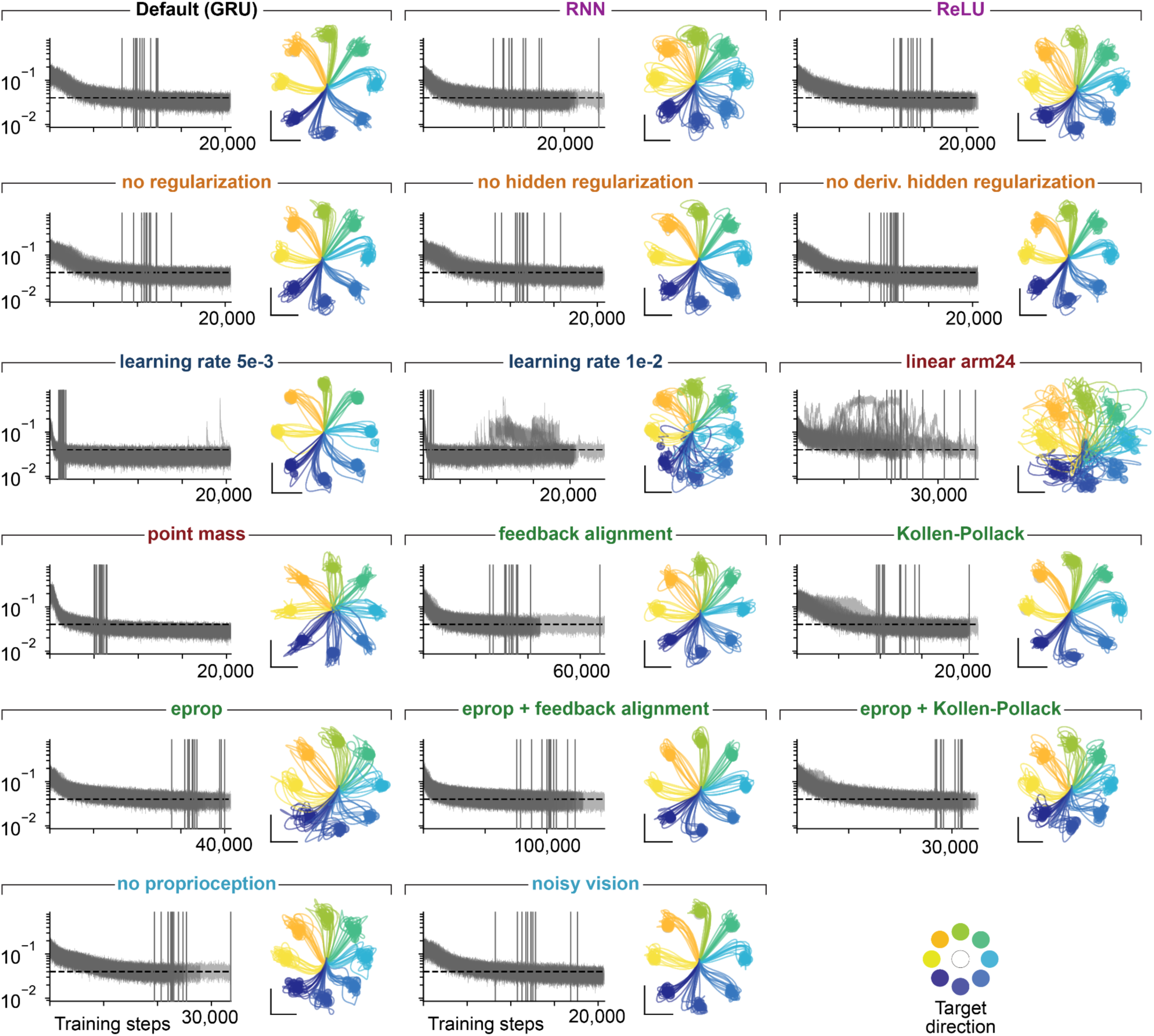
Training performance for all MotorNet architectural variations. Each subpanel shows one architectural variation in the same format. **Left:** the training loss 𝐿_!_ (excluding regularization terms) for the 12 seeds of a model variant. The black dashed horizontal line indicates the convergence threshold of 0.04 (see Methods), and the solid vertical lines indicate the first time a given seed met the convergence criterion of being under the threshold for 20 training steps on average. Note that some models kept training despite reaching the convergence criterion early as they had not reached the minimum number of training steps (20,480) required. **Right:** Endpoint kinematics produced for 8 center-out reach directions for all trained RNN architectural variations. The black vertical and horizontal lines indicate 5 cm lengths for the x and y dimensions.

**Figure S10.**
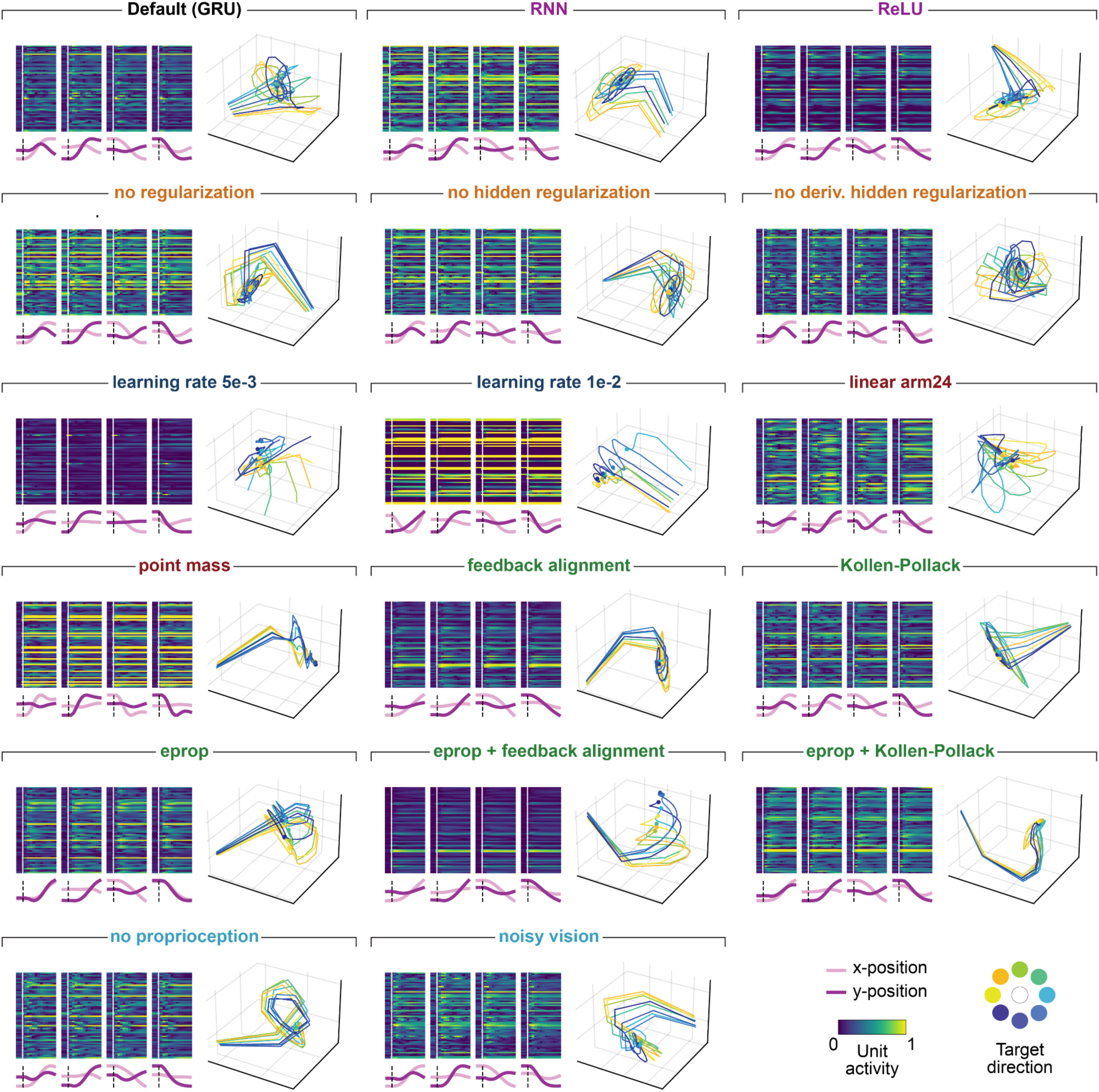
Example unit activations for all MotorNet architectural variations. Each subpanel shows hidden unit activations for four reach directions (left) and all example neural trajectories found by PCA (right; leading 3 neural modes) for all trained RNN architectural variants.

**Table S1.** *Test statistics and p-values*. All comparisons are Wilcoxon’s rank-sum tests.

| Dist. 1 | Dist. 2 | T. statistic | p-value |
| --- | --- | --- | --- |
| Fig. 3f |  |  |  |
| Ms w/in | Mk w/in | 2.062 | 0.039 |
|  | Hu w/in | 2.345 | 0.019 |
|  | Ms x Ms | -2.508 | 0.012 |
|  | Mk x Mk | -2.446 | 0.015 |
|  | Ms x Mk | -2.367 | 0.018 |
|  | Hu x Ms | -3.020 | 0.003 |
|  | Hu x Mk | -2.613 | 0.009 |
|  | Shuffle | -3.409 | 0.001 |
| Mk w/in | Hu w/in | 1.030 | 0.303 |
|  | Ms x Ms | -3.941 | 8.114e-5 |
|  | Mk x Mk | -4.440 | 8.984e-6 |
|  | Ms x Mk | -4.671 | 3.005e-6 |
|  | Hu x Ms | -4.752 | 2.017e-6 |
|  | Hu x Mk | -4.26 | 3.727e-6 |
|  | Shuffle | -5.760 | 8.398e-9 |
| Hu w/in | Ms x Ms | -3.501 | 4.640e-4 |
|  | Mk x Mk | -3.698 | 2.174e-4 |
|  | Ms x Mk | -3.853 | 1.168e-4 |
|  | Hu x Ms | -3.750 | 1.771e-4 |
|  | Hu x Mk | -3.784 | 1.546e-4 |
|  | Shuffle | -4.149 | 3.342e-5 |
| Ms x Ms | Mk x Mk | -2.672 | 0.008 |
|  | Ms x Mk | -0.171 | 0.864 |
|  | Hu x Ms | -2.435 | 0.015 |
|  | Hu x Mk | -2.494 | 0.013 |
|  | Shuffle | -6.241 | 4.351e-10 |
| Mk x Mk | Ms x Mk | 2.905 | 0.004 |
|  | Hu x Ms | -0.565 | 0.573 |
|  | Hu x Mk | -1.252 | 0.210 |
|  | Shuffle | -6.148 | 7.85e-10 |
| Ms x Mk | Hu x Ms | -3.326 | 8.795e-4 |
|  | Hu x Mk | -3.666 | 2.463e-4 |
|  | Shuffle | -10.42 | 1.919e-25 |
| Hu x Ms | Hu x Mk | -0.984 | 0.325 |
|  | Shuffle | -4.319 | 1.565e-5 |
| Hu x Mk | Shuffle | -3.147 | 0.002 |
| Fig. 4c |  |  |  |
| Shuffle | Somat. cortex | 2.422 | 0.015 |
|  | x regions (w/in) | -3.489 | 4.852e-4 |
|  | Motor Hu x Ms | 4.319 | 1.565e-5 |
|  | Motor Hu x Mk | 3.147 | 0.002 |
|  | x regions Hu x Ms | -6.846 | 7.603e-12 |
|  | x regions Hu x Mk | -7.996 | 1.289e-15 |
| Somatosensory cortex | x regions (w/in) | -2.997 | 0.003 |
|  | Motor Hu x Ms | -1.175 | 0.240 |
|  | Motor Hu x Mk | -1.924 | 0.054 |
|  | x regions Hu x Ms | -3.795 | 1.478e-4 |
|  | x regions Hu x Mk | -3.914 | 9.079e-5 |
| Across regions (w/in) | Motor Hu x Ms | 4.398 | 1.095e-5 |
|  | Motor Hu x Mk | 3.909 | 9.254e-5 |
|  | x regions Hu x Ms | 2.656 | 0.008 |
|  | x regions Hu x Mk | -2.932 | 0.003 |
| Motor Hu x Ms | Motor Hu x Mk | -0.984 | 0.325 |
|  | x regions Hu x Ms | -6.375 | 1.836e-10 |
|  | x regions Hu x Mk | -6.854 | 7.201e-12 |
| Motor Hu x Mk | x regions Hu x Ms | -6.729 | 1.708e-11 |
|  | x regions Hu x Mk | -7.352 | 1.959e-13 |
| x regions Hu x Ms | x regions Hu x Mk | -0.486 | 0.627 |
| Fig. 4e |  |  |  |
| Shuffle | x beh. w/in Mk | 0.564 | 0.573 |
|  | x beh x Mk | -1.616 | 0.106 |
|  | Exec. Mk x Ms | 10.424 | 1.919e-25 |
|  | Exec Mk x Hu | 3.147 | 0.002 |
| Across behaviors w/in Mk | x beh x Mk | -1.026 | 0.305 |
|  | Exec. Mk x Ms | 3.169 | 0.305 |
|  | Exec Mk x Hu | 1.92 | 0.091 |
| Across behaviors across Mk | Exec. Mk x Ms | 4.084 | 4.432e-5 |
|  | Exec Mk x Hu | 3.852 | 1.117e-4 |
| Exec. Mk x Ms | Exec Mk x Hu | -3.666 | 2.463e-4 |
| Fig. 5g |  |  |  |
| Ms x Ms | Mk x Mk | 5.745 | 9.216e-9 |
|  | Ms x Mk | 3.939 | 8.171e-5 |
|  | Hu x Ms | 5.572 | 2.523e-8 |
|  | Hu x Mk | 5.931 | 9.216e-9 |
| Mk x Mk | Ms x Mk | -8.029 | 9.850e-16 |
|  | Hu x Ms | 3.821 | 1.330e-4 |
|  | Hu x Mk | 3.802 | 1.437e-4 |
| Ms x Mk | Hu x Ms | 7.707 | 1.291e-14 |
|  | Hu x Mk | 8.631 | 6.091e-18 |
| Hu x Ms | Hu x Mk | 1.134 | 0.257 |

**Table S2.** *Summary of architectural and training variations explored with the MotorNet model*. When appropriate, acronyms used throughout the paper (e.g. **Fig. 6**) are defined in parentheses.

| Manipulation type | Default | Number of variations | Variations |
| --- | --- | --- | --- |
| None |  | 1 | Default model with all values in second column |
| Network architecture | GRU with tanh non-linearity | 2 | Vanilla recurrent units (RNN) |
|  |  |  | ReLU non-linearity (ReLU) |
| Training loss | Full regularization | 3 | No regularization of network hidden activity (no hidden reg.) |
|  |  |  | No regularization of the derivative of network hidden activity |
|  |  |  | No regularization in hidden activity or its derivative |
| Learning rate | 3e-4 | 2 | 5e-3 |
|  |  |  | 1e-2 |
| Learning algorithm | Backpropagation through time | 5 | Kollen-Pollack (K-P) |
|  |  |  | Feedback alignment (fb. alignment) |
|  |  |  | e-prop |
|  |  |  | e-prop + Feedback alignment |
|  |  |  | e-prop+ Kollen-Pollack |
| Inputs | Full sensorimotor feedback | 2 | No proprioception |
|  |  |  | Noisy vision |
| Environment | Two-joint arm with six Hill-type actuators | 2 | two-joints arms with two antagonist positive torque-producing actuators per joint (linear arm24) |
|  |  |  | Point mass |
| Total |  | 17 model variants |  |

